# Predicting Human Pharmacokinetic Parameters of Drugs using a Multi-Tissue Chip Platform Integrating Liver, Kidney, and Skeletal Muscle Microphysiological Systems

**DOI:** 10.64898/2025.12.19.695548

**Authors:** Jason Sherfey, Shiny Amala Priya Rajan, Lauren Nichol, Paarth Parekh, J. Tyler Smith, Lauren Gregory, Frances Clark, Shivam Ohri, Eugene P. Kadar, Billy T. George, David Tess, James R. Gosset, Jennifer Liras, Emily Geishecker, R. Scott Obach, Murat Cirit

**Affiliations:** Javelin Biotech Inc., 900 Middlesex Turnpike, Billerica, Massachusetts 01821, United States; Pfizer Global Research and Development, Groton Laboratories, Eastern Point Road, Groton, Connecticut 06340, United States; Pfizer Worldwide Research and Development, 610 Main Street, Cambridge, Massachusetts 02139, United States

**Keywords:** Microphysiological Systems, Tissue Chips, NAMs, pharmacokinetics, in vitro in vivo extrapolation

## Abstract

Accurate prediction of human pharmacokinetic (PK) parameters remains a critical challenge in drug discovery. While simple *in vitro* assays provide insight into discrete metabolic processes, they cannot recapitulate the complex multi-organ interactions that involve absorption, distribution, metabolism, and excretion (ADME) processes. This limitation has perpetuated a heavy reliance on animal experimentation to predict integrated outcomes like hepatic and renal clearance in humans. In this study, we developed a multi-tissue chip (MTC) to evaluate the PK parameters of intravenously (IV) administered drugs. The MTC platform provides continuous oxygenation through media recirculation and interconnects three MPSs (liver, kidney, skeletal muscle) to study drug metabolism, excretion and distribution within a single *in vitro* system. The functionality of each MPS was characterized for more than 15 days with single- and multi-tissue chips. A diverse set of drugs from all extended clearance classification system (ECCS) classes with various clearance mechanisms was evaluated on the MTCs and on-chip PK parameters were evaluated, such as hepatic metabolism, uptake, and disposition, tubular secretion and reabsorption, and muscle distribution. These PK parameters were successfully correlated to clinical parameters for IV drugs: hepatic clearance (CL_H_), renal clearance (CL_R_), volume of distribution (VD) and then these parameters were extrapolated to human drug exposures using physiologically-based pharmacokinetic (PBPK) modeling. This study demonstrated that the MTC platform could deliver accurate prediction for clinical pharmacokinetics of intravenously administered small molecules without the need for animal studies.

## Introduction

The current drug discovery paradigm faces a significant translational gap between preclinical findings and human clinical outcomes. Extrapolating pharmacokinetic (PK) parameters from traditional preclinical models, such as *in vitro* cell culture systems and animal studies, often proves unreliable due to inherent species-specific differences and limited predictability of human responses (Shanks N, 2009; Martignoni, 2006; H., 1995; Bowman, 2019). This challenge becomes more pronounced as new modalities, such as biologics, gene therapies, and cell-based therapies, emerge, creating further complexities like multi-tissue biodistribution patterns and requires human specific receptor expression and immune response that conventional models struggle to address (Benamara S, 2025; Deng, 2011; Xu X, 2012). Current traditional *in vitro* models, with their limited complexity and inability to replicate the multifaceted ADME (absorption, distribution, metabolism, and excretion) processes, fail to capture the intricate biological multi-organ interactions that are critical for drug development and clinical prediction (Godoy P, 2013; Fowler, 2020; Chunduri V, 2022). Additionally, the poor translation from animal data to clinical outcomes not only leads to the costly late-stage attrition of drug candidates but also raises significant ethical concerns regarding animal welfare, creating a strong imperative to adopt the principles of the 3Rs (Replacement, Reduction, and Refinement). To overcome these hurdles, there is a critical and growing need for New Approach Methodologies (NAMs) that can provide more predictive, human-relevant data early in the development pipeline (J., 2023; Sager JE, 2015; Fowler, 2020; Kang, 2024).

To bridge these gaps, advanced microphysiological systems (MPS), or “tissue chips,” have emerged as an innovative and more physiologically relevant alternative to traditional preclinical models. MPS platforms, particularly multi-tissue chips (MTCs), offer significant promise in mimicking human tissue microenvironments, and allowing simultaneous assessment of ADME parameters and their interdependencies to provide more accurate predictions of drug behavior in humans (Herland A, 2020; Edington CD, 2018; Tsamandouras, 2017; Abaci, 2015; Vernetti L, 2017). While MTCs have been increasingly recognized for their potential in drug development, most existing systems have been primarily designed for biological basic research applications and have yet to be fully optimized for pharmacokinetic (PK) studies.

Multi-tissue chip systems are complex *in vitro* platforms that integrate two or more tissues fluidically either in a flow-through format or via recirculating the basal medium. While flow-through multi-tissue chip systems can capture sequential tissue interactions, they miss the feedback loop of inter-organ communication and dynamic interactions of drugs and metabolites (Herland A, 2020; Vernetti L, 2017). As shown in our previous work (Rajan, et al., 2023), flow-through systems have major limitations in evaluating small molecules with a broad clearance range, particularly those with low turnover. In contrast, MTC systems with closed-loop recirculation allow dynamic drug interactions over time, mimicking *in vivo* conditions and allowing valuable PK data to be generated. However, current use of multi-MPS models in ADME face several limitations (Fowler, 2020): (i) they are often fabricated using cell lines that lack the necessary metabolic activity essential for accurate PK prediction; (ii) they have small tissue and medium volumes that become challenging during bioanalysis; (iii) they involve short-term integrated cultures with limited drug incubation times, making it difficult to evaluate low-clearance drugs and their metabolites; (iv) complex system setup that requires specialized expertise leading to slower adoption; (v) limited platform robustness causing frequent technical failures and reproducibility challenge and finally, (vi) they have not been adequately characterized for ADME applications.

Here, we developed and applied a human-relevant MTC platform that overcomes the limitations of traditional models by providing a robust and comprehensive platform for predicting multiple human PK parameters. Our system integrates three key ADME-relevant MPS, liver, kidney, and muscle that together enable a more holistic approach to studying drug metabolism, clearance, and distribution respectively. The liver MPS assesses hepatic clearance, the kidney MPS evaluates tubular secretion and reabsorption, and the muscle MPS investigates drug disposition. Each tissue component was characterized for functionality and metabolic activity over 15 days to ensure physiologically relevant conditions for PK studies. Novel computational methods were developed for the platform to translate time-concentration measurements from multiple compartments into clinical PK parameters. Here, we used this platform to evaluate a diverse set of small molecule drugs representing all Extended Clearance Classification System (ECCS) classes with a variety of clearance mechanisms, to estimate PK parameters from the MTC system, including hepatic clearance (CL_H_), renal clearance (CL_R_), and volume of distribution (VD). Our results show that the MTC accurately predicted these parameters, with a strong correlation to clinical data, highlighting its potential for improving the translation of preclinical data to human outcomes. In addition to its physiological relevance, this study incorporated complex physiologically based pharmacokinetic (PBPK) modeling, which allows for the additional prediction of human exposure and clinical PK profiles based on MTC-derived PK parameters.

## Materials & Methods

### Chip features and fabrication

The MTC was designed and fabricated from cyclic olefin copolymer (COC) thermoplastic, a transparent, biocompatible material that facilitates optical imaging while exhibiting minimal non-specific binding properties (Rajan, et al., 2023). Individual chip layers were precision-machined to incorporate three culture chambers and one oxygenation chamber, interconnected via microfluidic channels with 1 mm² cross-sectional areas, then assembled through thermal bonding. The culture chambers are enclosed using two distinct removable lid configurations: flat lids for direct tissue seeding within the chamber, and transwell adapter lids designed to integrate barrier tissues cultured on transwell inserts (**Fig. 1A**).

**Figure 1.**
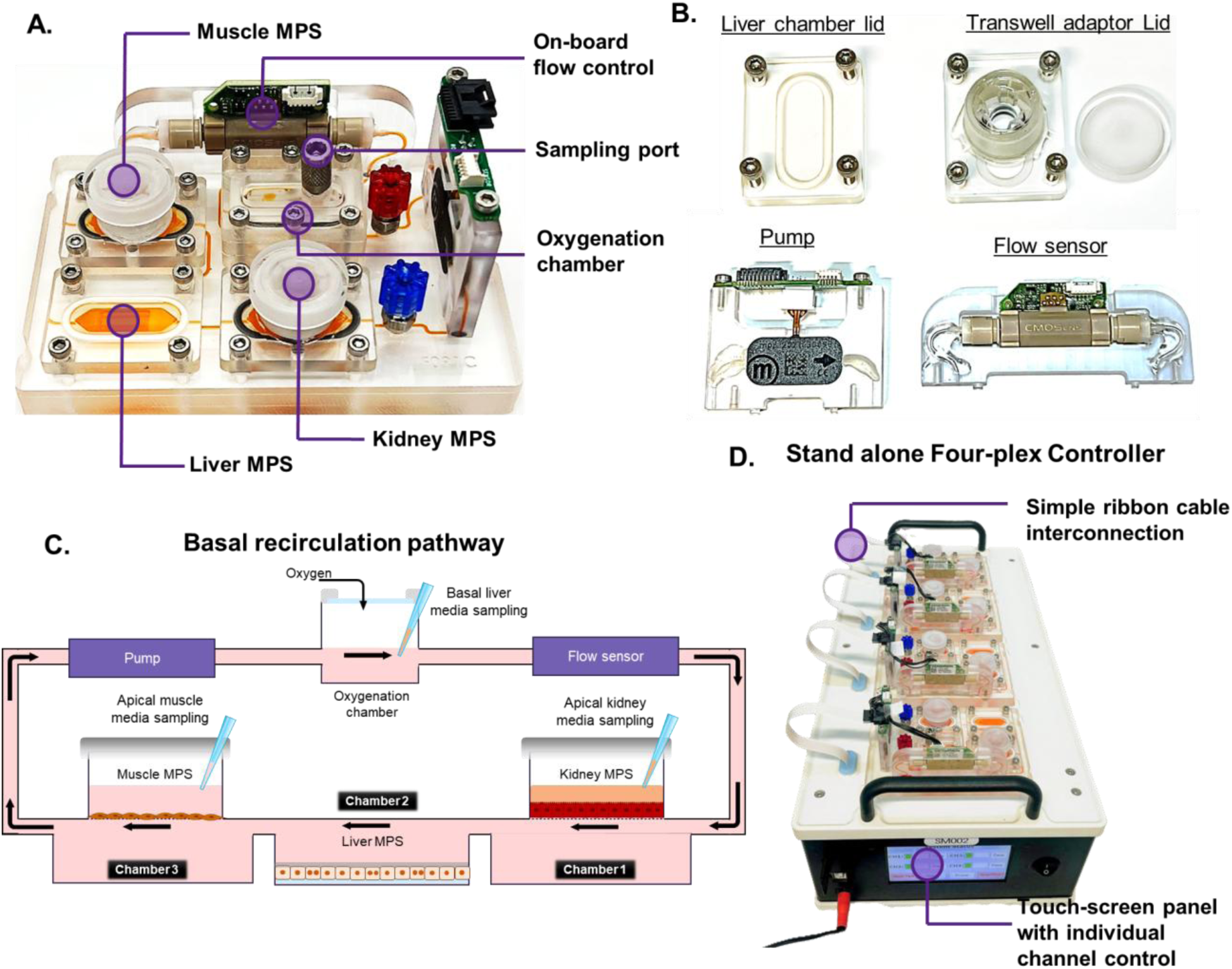
Overview of the Multi-Tissue Chip (MTC) system architecture and operational features. A) The MTC is designed with three separate tissue compartments housed in a single chip. B) These compartments are sealed either by liver chamber lid (Liver MPS) or an added transwell adapter for the kidney or muscle MPS. The system uses an on-board diaphragm pump and a flow sensor to maintain stable, recirculating basal flow that C) connects all three tissue chambers (liver, kidney, and muscle MPS). C) The additional oxygenation chamber maintains relevant oxygen levels in the media providing continuous oxygen to the MPS. This chamber also serves as a sampling port for basal medium throughout the experiment. The MTC media circulation is managed by D) stand-alone multiplexed controller, which is housed inside an incubator and features a user-friendly touch screen interface.

Key architectural features of the platform include (**Fig. 1A**): (i) a sampling port enabling easy accessibility for collecting kinetic media samples throughout experimental periods; (ii) removable tissue chamber lids providing direct access to cultured tissues during and end of study (**Fig. 1B**); (iii) on-board flow sensors coupled with piezoelectric pumps providing real-time flow monitoring and precise flow rate regulation (Fig. 1B); and (iv) an integrated oxygenation chamber facilitating continuous media oxygenation via recirculation (**Fig. 1C**); MTC along with the dual lid configuration provides flexible tissue integration options while maintaining controlled microenvironmental conditions essential for long-term culture stability (**Fig. 1B)**.

The basic working principle of the platform is similar to our previous publication on the liver tissue chip (Rajan, et al., 2023). Briefly, each chip features an integrated diaphragm pump that eliminates the need for external pumps, reservoirs, and tubing connections. This self-containing recirculating system enables dynamic media conditioning that promotes long-term culture while ensuring optimal oxygen and nutrient delivery to all the MPSs. The piezoelectric pumps accommodate a broad flow range (8 µL/min to >20 mL/min), allowing precise control of the media flow rate for various experiment conditions and tissue types. This chip also conforms to standard microtiter plate dimensions, ensuring compatibility with standard laboratory equipment.

A proprietary controller (Javelin Biotech Inc) operates up to four chips simultaneously **(Fig. 1D)**, featuring an intuitive interface for flow parameter configuration and real-time monitoring. This multiplexed design enhances experimental throughput, accommodating up to 24 chips within a single incubator.

### MPS cultures

#### Liver MPS

Cryopreserved primary human hepatocytes (PHH) donor lots were purchased from BioIVT (Donor Lot# WDH, Male (**Supp. Table 1**)). The liver MPS was seeded directly on the chip cell culture chamber and the DMPK performance and extensive characterization in a perfused sandwich culture format using this donor were previously reported using the Liver Tissue Chip (Rajan, et al., 2023).

Multi-tissue chip recirculation pathways and chambers were sterilized via 70% ethanol perfusion through all fluidic components (chambers, channels, and pump system), followed by 1-hour UV exposure of thermoplastic elements. Then, the cell culture chamber was pre-surface treated prior to extracellular matrix (ECM) coating. The PHH seeding protocol for MTC liver culture chamber (**Chamber 2, Fig. 1C**) followed established methods (Rajan, et al., 2023; Ohri, Parekh, Nichols, Rajan, & Cirit, 2024) The tissue chamber was coated with rat tail collagen I (100 μg/mL; Corning) and fibronectin (25 µg/mL; Sigma Aldrich) in cold PBS, then incubated overnight at 4°C. Prior to seeding, the chips were warmed to 37°C for at least 2 hours and washed with PBS or media.

The cryopreserved PHHs were then thawed and centrifuged at 100 × *g* for 8 min in pre-warmed cryopreserved hepatocyte recovery medium (CHRM, Gibco). The supernatant was aspirated, and cells re-suspended at 0.86 × 10^6^ cells/mL in pre-warmed hepatocyte plating medium (HPM), containing Williams’ E Medium (WEM) (Gibco) supplemented with primary hepatocyte thawing and plating supplements (Gibco). Each chip was seeded with 215,000 cells in 250 μL media and was incubated at 37°C in a 5% CO_2_ incubator for 4–6 h to allow cell attachment. Following incubation, the dead or unattached cells were removed from the surface by tapping the edge of the chip and aspirating the media. PHH culture was overlaid with 0.35 μg/mL Matrigel, (Corning) prepared in cold hepatocyte maintenance media (HMM; WEM plus hepatocyte maintenance supplements; Gibco). After overnight incubation, tissue chambers were sealed with a chamber lid and the chips were filled with 3 mL of HMM. Chips were connected to the controller installed in the incubator with the flow set at 2 mL/h.

#### Skeletal Muscle MPS

Primary human skeletal muscle myoblasts (HSMM; Lonza) were thawed and cultured off-chip on transwell inserts and matured in isolation. Briefly, HSMM were thawed, centrifuged for 5 min at 220 × *g*, and resuspended at 250,000 cells/mL in muscle growth medium (MGM) prepared by combining expansion supplements (STEMCELL Technologies; Myocult-SF Expansion Supplement Kit) with basal medium (DMEM with 1000 mg/L D-Glucose (STEMCELL Technologies). The HSMM cells were seeded on the apical side at 50,000 cells/cm2 on fibronectin-coated (10µg/cm2 bovine plasma fibronectin; Sigma,) transwell inserts (Corning). The cells were maintained in MGM both apically and basally overnight before replacing them with muscle differentiation medium (MDM; Human Myocult Differentiation Kit, STEMCELL Technologies) to induce myoblast differentiation into myotubes. The cultures were maintained at 37°C with media changes every 2–3 days until confluence reached 95-100% and multinucleated myotubes were visible throughout the culture.

#### Kidney MPS

Primary human Renal Proximal Tubule Epithelial Cells (RPTECs; ScienCell,) were expanded to passage 2 in a poly-L-lysine-coated culture vessel (2 μg/cm^2^; ScienCell) according to vendor instructions. Thawed cells were expanded in epithelial cell medium (EpiCM; ScienCell) supplemented with 2% FBS (ScienCell), 1% Epithelial Cell Growth Supplement (EpiCGS; ScienCell) and 1% penicillin/streptomycin solution (P/S; ScienCell). The cells were trypsinized from the vessel once the culture reached 90% confluence and were either cryopreserved or seeded directly to transwells (Corning) for off-chip maturation. Transwells were pre-coated with Human Collagen Type IV (100 µg/mL; Millipore Sigma) and fibronectin (25 µg/mL; Sigma Aldrich) and seeded at 75,000 cells/cm^2^. Cultures were maintained in EpiCM both apically and basally and changed every 2–3 days. Transepithelial electrical resistance (TEER) was monitored every 2 – 3 days to assess barrier integrity. Upon achieving a differentiated culture with hemicyst formation and a TEER ≥100 ohms.cm^2^, transwells were connected to the MTC via the adapter lid.

### MPS integration on the MTC

The MTC is a three-organ MPS platform integrating liver, kidney, and muscle tissues for PK studies **(Fig. 2A)**. Muscle and kidney MPSs were matured off-chip on transwell inserts as described above before integration into MTC. Kidney MPS was established first, as they require 8–10 days to mature and develop barrier integrity. Once kidney MPS TEER was ≥100 ohms.cm^2^, muscle MPS was seeded on the transwell at least 3 days before liver MPS culture. Upon reaching 95–100%confluency, muscle cultures were ready to be integrated to the MTC. Hepatocytes were directly seeded in the tissue chamber 2 as described above. Following the sealing of liver tissue chamber, 3 mL HMM was added and chips were connected to the controller and maintained under 2 mL/h flow. On day 3, the chamber 1 and chamber 3 lids were replaced with the kidney and muscle transwells via their respective adapter lid (**Fig. 1B**).

**Figure 2.**
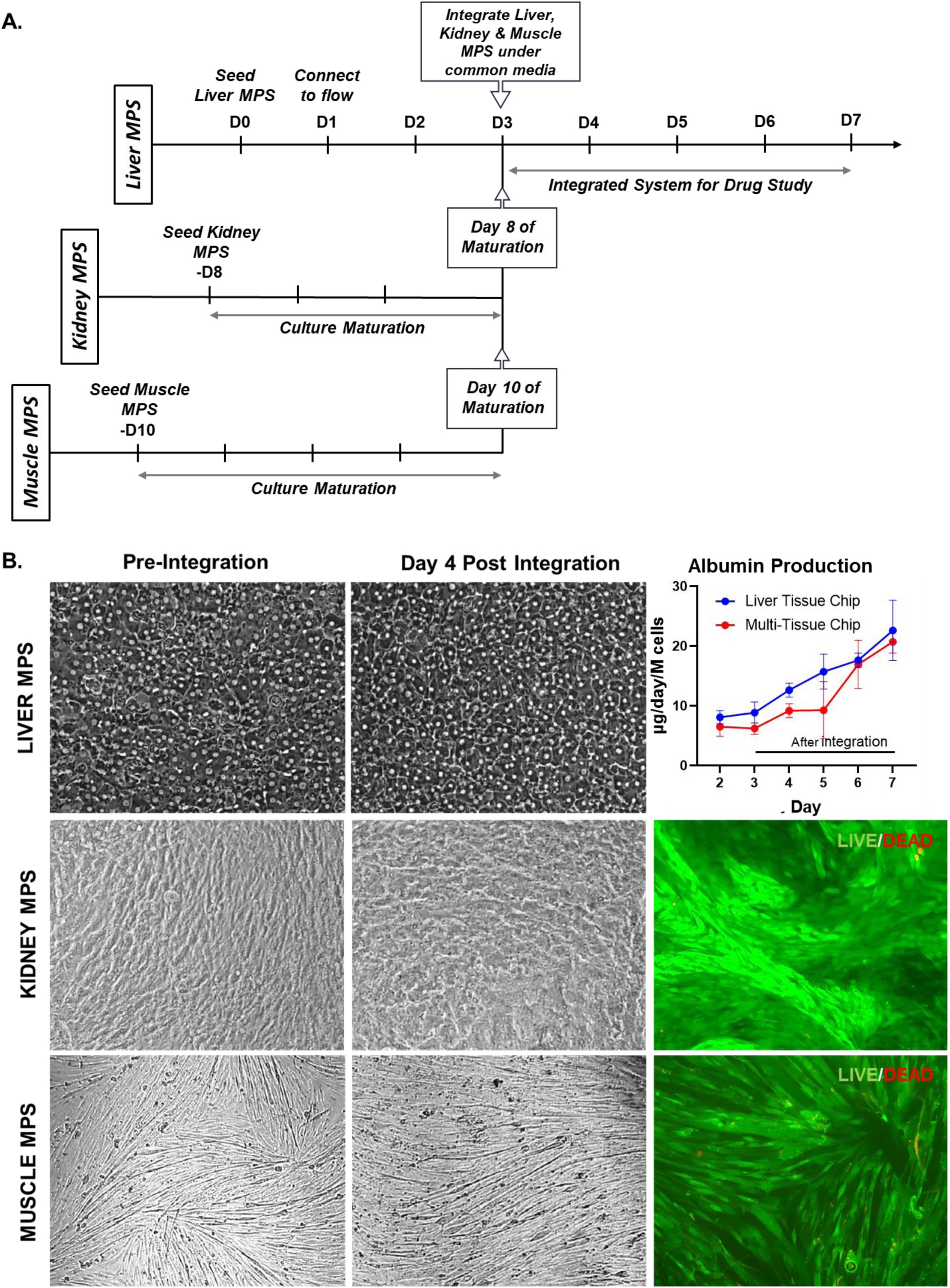
Integrated Multi-Tissue Chip (MTC) with three pharmacologically relevant tissues (liver, kidney and muscle) under common medium. A) The optimized experimental workflow for individual tissue culture maturation, integration, and maintenance in the MTC for PK drug studies. B) Individual tissue culture morphology before and after integration under common medium for 4 days. Albumin production is shown for liver functionality and live/dead staining for viability assessment of kidney and muscle.

The apical compartment of the kidney MPS was maintained in kidney-specific EpiCM medium, whereas the apical side of muscle and basal recirculation was maintained in common media prepared in HMM supplemented with 10% muscle growth supplement. Media were circulated in the integrated chip for 24 h before the drug study.

#### Common media for interconnected liver-kidney-muscle MPSs

To develop a common medium maintaining viability of all tissues while best preserving their physiology and functionality in the integrated MTC, we supplemented hepatocyte maintenance medium with growth factors from other tissue media as a strategy. Common media candidates were formulated by adding between 5% and 100% of the recommended quantities of muscle growth supplement (Stemcell Technologies; Myocult-SF Expansion Supplement Kit) and differentiation supplement (Myocult Differentiation Kit [Human], STEMCELL Technologies) to hepatocyte maintenance medium. Since the kidney MPS maintains barrier integrity with its maintenance medium (EpiCM; ScienCell) on the apical side, kidney supplements were excluded from the basal common medium. Common media candidates evaluated separately on muscle and liver cultures to identify an optimal formulation supporting both tissue types.

Individual MPS performance under common media was assessed in both isolated culture and in the interconnected platform with recirculating common media. Liver MPS evaluation included cell attachment efficiency, morphological analysis via brightfield imaging, and functional assessment through albumin production assays. Muscle MPS assessment focused on maintenance of differentiated morphology under brightfield microscopy and viability determination using LIVE/DEAD assay (**Supp. Fig. S2**). Kidney MPS, showing tight barrier functionality, was maintained apically with kidney-specific media, eliminating the need for common media compatibility assessment.

#### Phenotypic Characterization

For liver MPS, the functionality was assessed through quantification of albumin and urea production in culture supernatants using established immunoassay and colorimetric methodologies. Albumin concentrations were determined using the R-PLEX Human Albumin Antibody Set (Meso Scale Discovery) with media samples diluted 1:250 according to manufacturer protocols and analyzed on the MESO QuickPlex SQ120MM plate reader. Urea quantification employed the QuantiChrom Urea Assay Kit (BioAssay Systems) following vendor recommendations for low-concentration samples, with optical density measurements obtained at 430nm using a SpectraMax M3 Multi-mode Spectrophotometer. Production rates were normalized to cell number, chip volume, and media exchange parameters to yield daily albumin and urea production per million hepatocytes.

Bile canaliculi visualization was achieved through live cell staining using 5-chloromethylfluorescein diacetate (CMFDA, Invitrogen). Culture media was replaced with CMFDA-containing maintenance medium prepared by diluting 10mM stock solution 1:500. Following a two-hour incubation under continuous flow conditions, CMFDA medium was exchanged with fresh hepatocyte maintenance medium prior to imaging. Fluorescent microscopy was performed at 488nm excitation with simultaneous brightfield image acquisition to enable overlay visualization of bile canalicular networks within the hepatocyte monolayer.

#### Barrier Integrity Measurements

Barrier integrity of kidney MPS was assessed under static conditions by transferring transwell inserts from chip-mounted adaptors to standard 24-well plates or well plates associated with the transwells. Transepithelial electrical resistance (TEER) measurements were obtained using sterile, preconditioned STX2 electrodes (World Precision Instruments) connected to an EVOM2™ Epithelial Volt/Ohm Meter (World Precision Instruments) positioned according to manufacturer specifications. Stable resistance values were recorded and normalized to baseline measurements obtained from blank wells.

Permeability assessment employed fluorescently labeled tracers including 3000 MW dextran (Molecular probe dextran; Fisher Scientific) or Lucifer Yellow (LY; Sigma-Aldrich, Catalog# L0144) added to either apical or basolateral compartments of the kidney model. Tracer leakage from donor to receiver compartments was quantified through fluorescence spectrophotometry following two-hour incubation periods. The resultant leakage ratio served as a quantitative measure of renal proximal tubular epithelial cell (RPTEC) monolayer integrity.

#### Immunocytochemistry Staining

For IHC staining, the MPS was fixed with 4% paraformaldehyde for 15 min at room temperature (RT) or overnight at 4ᵒC and permeabilized by incubation in 0.2% Triton-X for 10 min at RT. Non-specific antibody binding was blocked by incubation in Protein Block Solution (Abcam) for 30 min in a rocker/shaker at RT. The primary antibodies diluted in antibody diluent (Agilent Technologies) were added and incubated overnight at 4ᵒC. The hepatocytes were stained with E-Cadherin (Abcam) MRP-2 (Fisher Scientific), OAT1 (Santa Cruz Biotechnology), OAT3 (Santa Cruz Biotechnology), zonula occludens-1 (ZO-1; Fisher Scientific,), Anti-Aquaporin 1 antibody (AQP-1; Abcam), and Myosin Heavy Chain (MHC; R&D Systems). Following primary incubation, the tissues were washed in PBS and incubated for 1 hr with anti-rabbit, anti-mouse, Alexa Fluor 488, 594 and 647 secondary antibodies (Invitrogen) as appropriate in antibody diluent (1:200 dilution). Cells were counterstained with DAPI for 7 min at RT and washed with PBS prior to fluorescent imaging. The tissue in the LTC or transwell was imaged using a Zeiss LSM880 upright confocal microscope or inverted epifluorescence microscope (Nikon Eclipse TE-2000S).

#### LIVE/DEAD cell viability

Cell viability was assessed by Live/Dead (L/D) assay. Briefly, the stain solution was made by preparing 2 μM calcein-AM and 2 μM ethidium homodimer-1 (Invitrogen, Live/dead™ Viability/Cytotoxicity Kit, for mammalian cells, Thermo Fisher) in a 1:1 mixture of media and PBS. MPS was washed with fresh PBS prior to adding the L/D solution into the culture chamber or transwell. After 15 min incubation, the MPS was washed again with PBS before imaging with inverted epifluorescence microscope (Nikon Eclipse TE-2000S). Representative images per MPS were obtained using 405 nm (green) and 559 nm (red) fluorescence filters for calcein-AM indicating live cells and ethidium homodimer indicating dead cells respectively.

#### Drug preparation

Drug stock solutions were prepared by dissolving powdered compounds in DMSO at concentrations minimally 1000-fold above target working concentrations and stored at -80°C until experimental use. Midazolam (Sigma-Aldrich) and alprazolam (Sigma-Aldrich) were obtained as pre-solubilized methanol solutions and subsequently diluted with DMSO to achieve 1mM stock concentrations on the day of study. Multi-drug cocktails were formulated by diluting individual stock solutions in culture media to achieve final concentrations of 1 µM (with the exception of propranolol at 0.1 µM) while maintaining DMSO content below 0.1%. Drug combinations were strategically designed to minimize potential drug-drug interactions (DDI) that could confound clearance predictions (**Supp. Table 2**). Specifically, compounds metabolized by identical cytochrome P450 isoforms were segregated into separate cocktails, as were substrates sharing common transporter pathways. Additionally, all drug combinations underwent validation for analytical compatibility to ensure reliable quantification via LC-MS/MS methodology within the same analytical mixture.

### Pharmacokinetic studies on MTC

Following 24 h of fluidic integration of liver, kidney, and muscle MPSs, the basal circulation media was completely replaced with 3mL of drug cocktail media (**Supp. Table 2**) prepared in common media formulation. Test compounds were added as a bolus dose at an initial concentration of 1 µM each, except propranolol at 0.1 µM, and the final solvent (dimethyl sulfoxide, DMSO) concentration never exceeded 0.1% (v/v). The muscle apical compartment received 250 µL of drug-containing media, while the apical kidney compartment received 250 µL of drug-free tissue-specific EpiCM medium. Media samples (25–50 µL) were collected from the reoxygenation chamber, the apical kidney compartment, and the apical muscle compartment at 1, 4, 8, 24, 48, and 72 h. At the end of the drug study, the muscle and liver MPSs were lysed with methanol for intracellular drug quantification. Drug concentrations were quantified using LC/MS and plotted as concentration-time profile to access drug depletion kinetics (**Supp. Fig. 3**). Detailed bioanalysis methodologies are provided in the Supplementary Methods.

**Figure 3.**
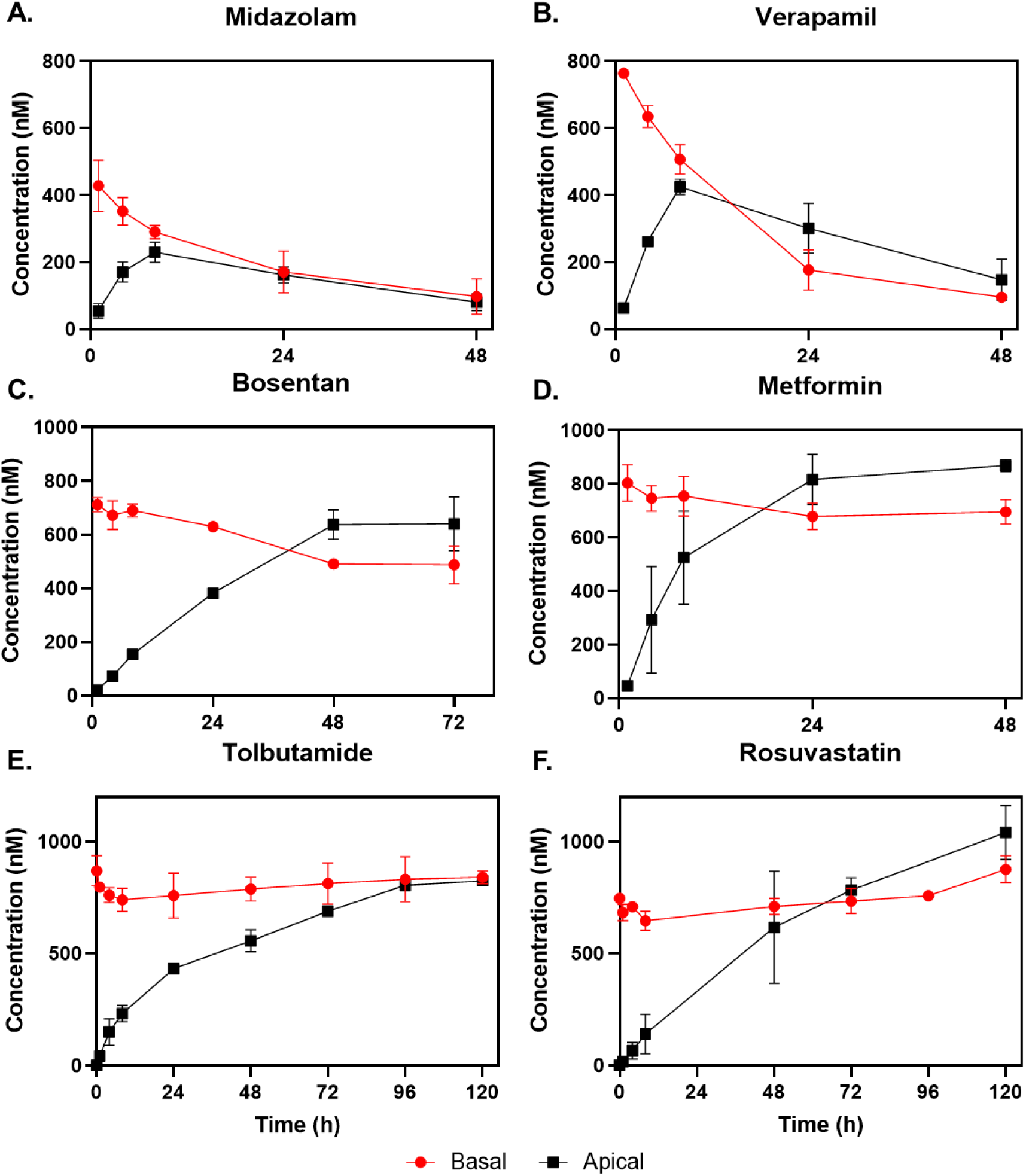
Representative drug study results from multi-tissue chips (MTCs) demonstrating different clearance mechanisms. A) Midazolam plot shows high hepatic metabolic clearance with no renal secretion. B) Verapamil shows mixed clearance through high passive transport and reabsorption from apical to basal kidney for metabolic clearance. C) Bosentan and D) Metformin have low metabolic clearance but are actively secreted to the apical kidney. E) Tolbutamide has low metabolic clearance and no renal secretion. F) Rosuvastatin has high biliary excretion, very low metabolic clearance, and high active renal secretion. Red lines show basal drug kinetics; black lines are apical kidney profiles.

### Computational data analysis for the MTC

The analysis and translation of MTC data involves: (1) *in vitro* data analysis to extract on-chip parameters for liver, kidney, and muscle; (2) translation of the on-chip parameters to predict *in vivo* PK parameters and PBPK concentration profiles. The MTC analysis assumes that the recirculating drug is metabolized in the basal liver compartment, transports across the kidney monolayer from basal to the apical kidney compartment, and finally the drug distributes into cells within each tissue compartment (muscle, liver, kidney). Kinetic media samples were collected both from apical and basal compartments throughout the study, and intracellular samples were obtained from cell lysates at the end of the experiment.

### *In vitro* data analysis

The *in vitro* data analysis involved extracting key PK parameters associated with each organ (liver, kidney, muscle) from kinetic drug concentration data obtained from the MTC study.

#### Liver MPS intrinsic clearance (CL_int,u_)

Intrinsic hepatic clearance was estimated from the depletion of the drug from the MTC using conservation of mass, non-compartmental analysis, and the inverse well-stirred model. First, conservation of mass was used to calculate the change in total drug (combined across compartments) from the sampled data. Assuming that loss of parent compound in the MTC was due to hepatic metabolism, the on-chip apparent clearance was estimated using noncompartmental analysis. Finally, the well-stirred model was applied to back-calculate the intrinsic hepatic clearance while accounting for protein binding and recirculating flow. See Supplementary Methods for equations and further details.

#### Kidney MPS apparent permeability (P_app, BA_ and P_app, AB_) and efflux ratio (ER_ss_)

The kidney MPS analysis involved using traditional *in vitro* apparent permeability calculation for P_app,BA_ and leveraging the steady state ER_ss_ observed in the long-term study to estimate P_app,AB_ from the same study. It was assumed that the quasi-steady state apical/basal concentration ratio in the kidney MPS approximates the efflux ratio. With this assumption, the permeability was estimated in both directions across the kidney monolayer from a study that includes dosing on the basal side without dosing on the apical side. See Supplementary Methods for further details.

#### Muscle MPS lysate/media ratio

On-chip disposition was quantified by the endpoint ratio of drug concentration in the muscle MPS lysate over the media sampled from the muscle MPS compartment. Based on an observed correlation, an empirical scaling factor was derived to translate this parameter to predict the *in vivo* steady state volume of distribution (VD_ss_) (see supplemental figures for data **(Supp. Fig. S5 – S8)**).

*In vitro in vivo* correlation (IVIVC) methods for translating the *in vitro* parameters (CL_int,u_, P_app,BA_, P_app,AB_, and lysate/media) to clinical PK endpoints (CL_H_, CL_R_, CL_T_, and VD_ss_) are detailed in the Supplementary Methods.

### *In vitro in vivo* extrapolation (IVIVE) and PBPK modeling

PK-Sim® (Open Systems Pharmacology Suite) (Willmann, 2003) was used for PBPK modeling. IVIVE using PBPK modeling was performed to predict human time-concentration profiles for representative drugs with different clearance mechanisms: midazolam (hepatically cleared), metformin (renally cleared), and verapamil (mixed clearance). Open-source models for each drug were adapted from the PK-Sim model library (Open systems pharmacology. GitHub. (n.d.). https://github.com/Open-Systems-Pharmacology) by replacing predefined metabolic processes with the MTC-derived human hepatic plasma clearance (CL_H_), representing an aggregate measure of the total impact of individual enzymes to drug metabolism. Similarly, predefined renal (e.g., glomerular filtration rate-GFR) processes in the PK-Sim model were replaced with the MTC-derived human renal clearance (CL_R_) predicted by IVIVC. AUC_0-inf_ values for predicted and clinically observed drug exposures were calculated using SimBiology (sbionca function) in MATLAB R2020a (MathWorks, Inc.). Detailed computational methodologies, including mathematical equations, step-by-step calculations, and model adaptations, are documented in Supplementary Information.

## Results

### Multi-Tissue Chip (MTC) platform for interconnecting three MPSs

The MTC platform enables interconnection of three MPSs, specifically liver-kidney-muscle MPS to evaluate metabolism, excretion, and disposition of drugs, respectively. The MTC is a millifluidic system that is fabricated used high-grade thermoplastic (COC) that has desirable properties like (i) high optical clarity that allow use of phase and fluorescence imaging to visualize cells, (ii) non-gas permeable material that makes the chip experiences a very low evaporation rate (< 0.56%/day (Rajan, et al., 2023)) and (iii) very low to negligible non-specific binding to most compounds (Rajan, et al., 2023) making it well-suited for PK applications. The MTC operates as a closed-loop recirculating system with controlled oxygenation to the culture, where ambient oxygen from the incubator diffuses through a gas-permeable silicone membrane overlying the oxygenation chamber as previously described (Rajan, et al., 2023). The integrated pumping system, featuring piezoelectric pumps and a flow sensor, enables dynamic media conditioning through continuous steady oxygenation in the chamber, which then recirculates through the MPS chamber. The reoxygenation chamber also acts like a bubble trapper to remove any bubbles that are formed in the fluidic system during the perfusion. The tissues in the MTC are interconnected through basal recirculation of the kidney, muscle, and over liver **(Fig. 1C)** for multi-tissue interaction, which are all regulated by the multiplex controller **(Fig. 1D)**. The platform enables comprehensive data acquisition through integrated imaging capabilities, kinetic media sampling, and endpoint tissue lysis.

### Assessment of the liver, kidney, and skeletal muscle MPS for viability and functionality

#### Liver MPS

The liver MPS was seeded into the MTC as a PHH sandwich culture to maintain physiological functionality and metabolic activity before and after integration with other MPSs. The liver MPS in this system was used to assess hepatic clearance of the drugs. By seeding PHH as a sandwich culture, hepatocyte morphology and liver-specific protein production (albumin, urea) were maintained for at least 15 days (Supp. **Fig. S1**). The polarity of the liver MPS was observed by IHC staining of bile canaliculi using CMFDA and MRP-2. The liver MPS maintained long-term CYP activity levels, as reported previously (3, 4)

#### Kidney MPS

The primary human RPTECs were cultured in a monolayer on semi-permeable transwell inserts to evaluate tubular secretion and reabsorption of drug molecules. RPTECs formed a uniform monolayer on the transwell membrane and differentiated to form tight junctions. During maturation, the culture developed dome-like structures or hemicyst, which are indicative of transporting epithelial cells (Van der Hauwaert C, 2013). The three dimensionality of these dome structures were evident in confocal images **(Supp Fig. S1)** of (ZO-1)-stained kidney tissue. Barrier function of the kidney tissue was investigated using TEER measurements and permeability assays. The TEER values were minimal at the initial days as the culture was reaching confluency and increased as the tissue was maturing to 100 – 200 Ω.cm^2^. Drug studies were conducted when the culture reached a TEER threshold of ≥100 Ω.cm^2^. The permeability assay with LY was performed under static conditions. Compared to the no-cell control group, only 6% of LY and 13.5% of dextran were transported through the tissue, showing great barrier function (Supp. **Fig. S1**).

Tight junction formation was visualized through ICC staining of the ZO-1 proteins which localize specifically to tight junction boundaries. Both dome-forming and monolayer cell populations demonstrated robust barrier integrity, with ZO-1 protein expression appropriately localized to cellular membranes. The water channel aquaporin-1 (AQP1) was expressed widely in plasma membranes in epithelial cells of the proximal tubule, confirming cellular differentiation and functional maturation. ICC staining revealed positive expression of organic anion transporters, OAT1 and OAT3, characterized by multiple fluorescent spots and diffused labeling in the cytoplasm. These transporters represent critical uptake mechanisms predominantly localized to basolateral membranes of RPTECs (5) and serve as primary mediators for drug secretion or transport from systemic circulation into urine (6,7). The presence and polarized localization of these transporters confirmed the functional relevance of the model for renal drug disposition studies.

#### Skeletal muscle MPS

The muscle MPS was used to quantify the tissue-plasma partition coefficient (K_p_), which can be extrapolated to the volume of distribution at steady state (VD_ss_) of a drug in humans. The muscle MPS was cultured using primary HSMM on Transwell inserts. The confluent monolayer of HSMM matured and differentiated to form multinucleated myotubes, indicating good culture health and differentiated morphology. The muscle MPS was stained for myogenic and myotopic markers to determine duration of myotube formation in cultures and myosin heavy chain (MHC) was expressed on this culture from day 5 onwards indicating differentiated and matured muscle culture **(Supp. Fig. S1).**

### MTC integration of liver, kidney, and skeletal muscle MPSs and functional assessment for PK studies

While single tissue chips can be used to generate MPS-specific data and to test tissue-specific hypotheses, MTCs provide a unique platform to study parent drugs and their metabolite kinetics simultaneously with multiple ADME-relevant tissues. Hence, we developed an integrated system for IV drug testing consisting of liver, kidney, and muscle MPS that maintain their functions in common media. This requires a systematically developed common medium that not only can support MPS viability but also maintain tissue functionality and phenotype relevant for PK application **(Supp. Fig. 2).**

In the MTC, the liver MPS was directly seeded on the chip, but muscle and kidney MPSs were matured off-chip on a Transwell as each tissue has different maturity timeframe (**Fig. 2A**). During integration, the kidney MPS with a tight barrier is maintained apically by the kidney-specific media. However, as muscle has a leaky barrier, the muscle-specific media will interact with the liver media. This makes it critical to develop a common media that both supports the functionality and enzyme activity of the liver and the differentiated phenotype of the muscle. HMM was selected as the base media to preserve the liver functionality that supports drug metabolism. To support the differentiated phenotype of the muscle, different concentrations of the muscle differentiation supplement (100%, 75%, 50%, 25%) and growth supplement (10%) were added to HMM and tested. Morphological and viability data suggested that after the maturation period, muscle cells could be maintained in HMM media containing <25% muscle differentiation supplement **(Supp. Fig. S2A).**

Next, the effect of muscle supplements in the common media on liver MPS was tested. To minimally influence the PHH with the muscle supplement, a range between 5% and 25% was tested. The cultures were incubated for 6 days, during which samples were collected for albumin and urea assays and assessment of CYP activity levels. As seen in **Supp. Figs. 2B–D**, there was no significant difference in albumin production between the control and the 5% differentiation supplement group but 15 – 20% decline is observed for the 12.5% and 25% group differentiation supplement group. However, the liver MPS functionality deteriorated in the muscle growth supplement group, as CYP activity decreased by 80% and albumin production by 72%. Based on these data, we concluded that any concentration of muscle growth supplement might deteriorate liver tissue functionality and to maintain healthy liver MPS, a common media with 5-10% muscle differentiation supplement could be used.

For PK studies on MTC, it is essential that the liver MPS matures with sustained metabolic and transporter activities, the kidney MPS has barrier integrity and maintains transporter activity, and the muscle MPS is fully differentiated. To achieve this, each tissue was matured in their respective preferred media before being integrated in the common medium. The liver MPS was mature for 3-4 days, the muscle tissue took up to 5 days to become fully confluent and for myoblasts to fuse to form myotubes and multinucleated cells, and the kidney took between 12 – 20 days to develop tight junctions with a TEER value >100 ohms.cm^2^. These are the entrance criteria used to identify tissues that can be integrated for drug studies.

The liver MPS had no morphological differences after 4 days of integration and the albumin production pattern was comparable to the control in a liver tissue chip **(Fig. 2B)**. The muscle tissue maintained its confluency and differentiated phenotype with high viability. Similarly, the kidney tissue was able to maintain its barrier integrity with a TEER value >100 ohms.cm^2^ with high viability. This demonstrated our ability to integrate three different MPS under common recirculating media and maintain individual organ functionality and phenotype to run PK studies.

### Assessment of PK parameters for IV drugs on MTC

The matured MPSs were maintained under common media for at least 24 h before the drug cocktails were introduced into the MTC. Nineteen drugs (dextromethorphan, diclofenac, midazolam, propranolol, alprazolam, tolbutamide, bosentan, S-warfarin, theophylline, zidovudine, fluvastatin, pitavastatin, raloxifene, verapamil, repaglinide, rosuvastatin, desipramine, metformin, penicillin G) were cocktailed in six groups (**Supp. Table 1**) and added to the basal recirculation compartment and apical muscle. No drug was added to the apical side of kidney to quantify drug secretion. Samples were taken both basally and apically at 1, 4, 8, 24, 48, 72, 96, and 120 hours.

The estimated PK parameters from the MTC included parent drug depletion in the basal side due to hepatic metabolism, muscle disposition, and renal clearance by secretion to the kidney apical side. In the MTC, 19 drugs were added as cocktails of multiple drugs that do not create drug-drug interaction issues. Midazolam is a high clearance drug that was nearly depleted in the basal compartment within 48 h; it permeated the kidney tissue with no apparent active transport since there was no concentration differential between the apical and basal sides. The predominant clearance mechanism of midazolam in the chip was hepatic clearance (**Fig. 3A**) which is consistent with what is known for in vivo human disposition of this drug. Verapamil showed mixed clearance through high passive transport and reabsorption from apical to basal kidney for metabolic clearance (**Fig. 3B**). In contrast, bosentan had low metabolic clearance but was actively secreted to apical kidney (**Fig. 3C**). Metformin was renally cleared, eliminated by tubular secretion by hOCT2 transporters localized to the basolateral side. It had minimal hepatic clearance and low passive permeability, which was observed as the parent drug concentration increases apically through renal uptake (**Fig. 3D**). There was also polarization of the kidney tissue in the MTC. In contrast, the low-clearance drug tolbutamide took more than 120 hours to observe at least 30% drug depletion, but the kidney accumulation kinetics suggested no active tubular transport (**Fig. 3E**). Rosuvastatin, on the other hand, was renally cleared, especially by tubular secretion involving BCRP, OAT1, and OAT3, the active transporters localized to the basolateral side. However, it had minimal hepatic clearance and low passive permeability but was actively secreted to the apical kidney (**Fig. 3F**).

### IVIVC and IVIVE of PK parameters from the MTC

A diverse set of drugs (**Supp. Table 1**) representing all Extended Clearance Classification System (ECCS) classes, with various clearance mechanisms, was tested on the MTC. The on-chip parameters (CL_int,u_, P_app, BA_, P_app, AB_, muscle lysate/media) necessary for downstream clinical PK predictions were calculated. The MTC was assessed across four key PK parameters—renal clearance (CL_R_), hepatic clearance (CL_H_), total clearance (CL_T_), and volume of distribution (VD_ss_)— by comparing MTC-based predictions to the corresponding clinical values.

Hepatic clearance, as shown in **Table 1** and **Fig. 4A**, was predicted well by the MTC model for all the ECCS classes. On-chip intrinsic hepatic clearances observed on the MTC, ranging 3.01 μl/min/million cells (rosuvastatin) to 321 μl/min/million cells (diclofenac), were scaled to predict *in vivo* hepatic clearances (in the Supplementary Materials). The predicted human (*in vivo*) hepatic clearance values spanned 0.26 ml/min/kg for alprazolam to 19.8 ml/min/kg for zidovudine. The MTC predicted hepatic clearance was within the three-fold error line for drugs like midazolam, diclofenac, fluvastatin, tolbutamide, and repaglinide, where hepatic clearance is the predominant route of elimination. The IVIVC of these drugs showed that >30% of drugs were within a two-fold error and >60% were within a three-fold error. The average fold error for all the drugs was 4 for the predicted hepatic clearance (ml/min/kg).

**Table 1.**
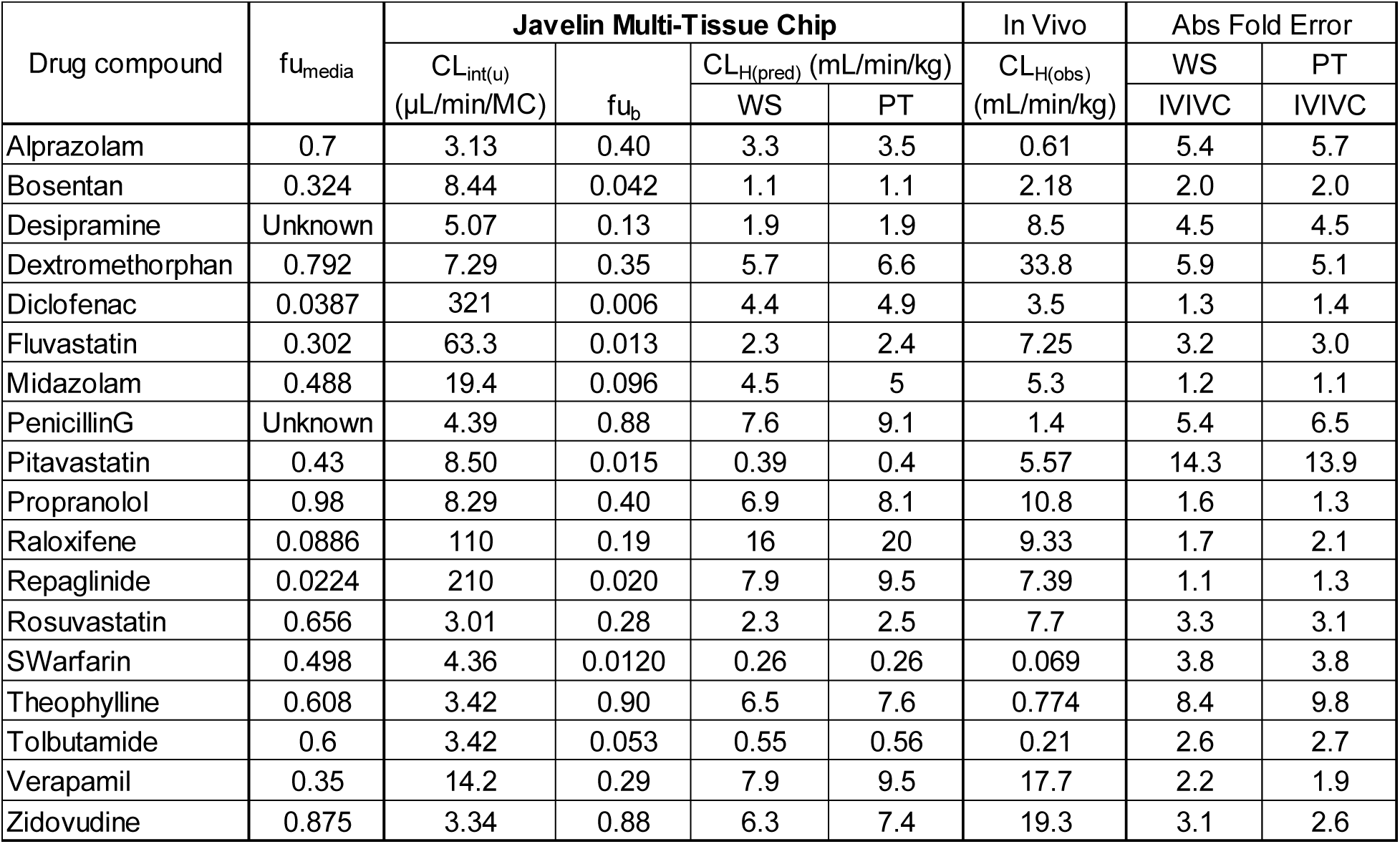
*In vitro in vivo* correlation (IVIVC) for hepatic clearance.

| Drug compound | fu<sub>media</sub> | Javelin Multi-Tissue Chip |  |  |  | In Vivo | Abs Fold Error |  |
| --- | --- | --- | --- | --- | --- | --- | --- | --- |
|  |  | CL<sub>int(u)</sub><br>(μL/min/MC) | fu<sub>b</sub> | CL<sub>H(pred)</sub> (mL/min/kg) |  | CL<sub>H(obs)</sub><br>(mL/min/kg) | WS | PT |
|  |  |  |  | WS | PT |  | IVIVC | IVIVC |
| Alprazolam | 0.7 | 3.13 | 0.40 | 3.3 | 3.5 | 0.61 | 5.4 | 5.7 |
| Bosentan | 0.324 | 8.44 | 0.042 | 1.1 | 1.1 | 2.18 | 2.0 | 2.0 |
| Desipramine | Unknown | 5.07 | 0.13 | 1.9 | 1.9 | 8.5 | 4.5 | 4.5 |
| Dextromethorphan | 0.792 | 7.29 | 0.35 | 5.7 | 6.6 | 33.8 | 5.9 | 5.1 |
| Diclofenac | 0.0387 | 321 | 0.006 | 4.4 | 4.9 | 3.5 | 1.3 | 1.4 |
| Fluvastatin | 0.302 | 63.3 | 0.013 | 2.3 | 2.4 | 7.25 | 3.2 | 3.0 |
| Midazolam | 0.488 | 19.4 | 0.096 | 4.5 | 5 | 5.3 | 1.2 | 1.1 |
| PenicillinG | Unknown | 4.39 | 0.88 | 7.6 | 9.1 | 1.4 | 5.4 | 6.5 |
| Pitavastatin | 0.43 | 8.50 | 0.015 | 0.39 | 0.4 | 5.57 | 14.3 | 13.9 |
| Propranolol | 0.98 | 8.29 | 0.40 | 6.9 | 8.1 | 10.8 | 1.6 | 1.3 |
| Raloxifene | 0.0886 | 110 | 0.19 | 16 | 20 | 9.33 | 1.7 | 2.1 |
| Repaglinide | 0.0224 | 210 | 0.020 | 7.9 | 9.5 | 7.39 | 1.1 | 1.3 |
| Rosuvastatin | 0.656 | 3.01 | 0.28 | 2.3 | 2.5 | 7.7 | 3.3 | 3.1 |
| SWarfarin | 0.498 | 4.36 | 0.0120 | 0.26 | 0.26 | 0.069 | 3.8 | 3.8 |
| Theophylline | 0.608 | 3.42 | 0.90 | 6.5 | 7.6 | 0.774 | 8.4 | 9.8 |
| Tolbutamide | 0.6 | 3.42 | 0.053 | 0.55 | 0.56 | 0.21 | 2.6 | 2.7 |
| Verapamil | 0.35 | 14.2 | 0.29 | 7.9 | 9.5 | 17.7 | 2.2 | 1.9 |
| Zidovudine | 0.875 | 3.34 | 0.88 | 6.3 | 7.4 | 19.3 | 3.1 | 2.6 |

**Figure 4.**
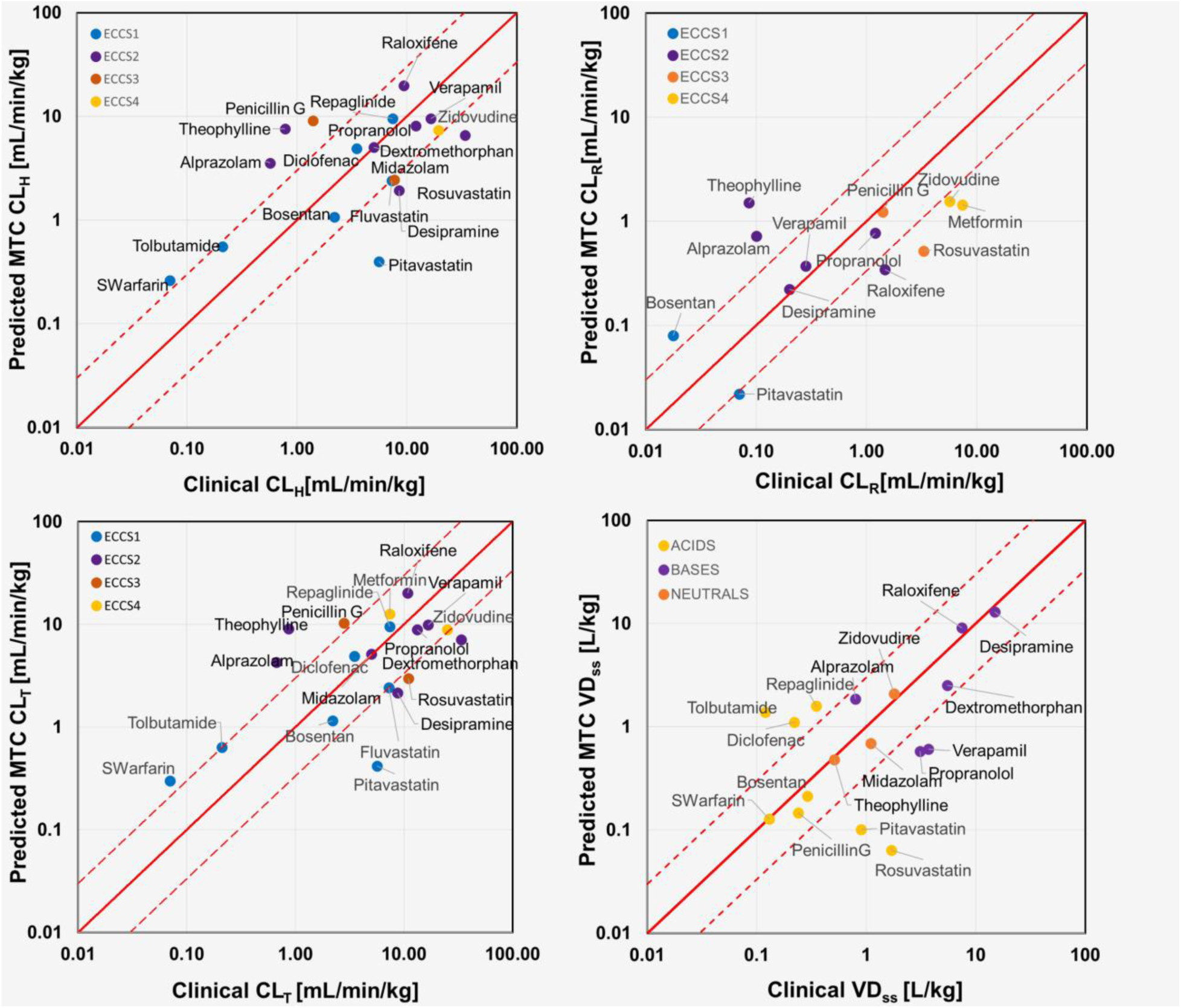
In vitro in vivo correlation (IVIVC) plots for PK endpoints predicted from a single MTC study: (A) hepatic clearance, (B) renal clearance, (C) total clearance, (D) steady state volume of distribution. Dashed lines mark three-fold difference between MTC predicted and clinically observed values.

Renal clearance predictions, as seen in **Table 2** and **Fig. 4B**, were accurate for most drugs, especially for those where renal excretion is the predominant route of elimination. On-chip kidney permeabilities (P_app, BA_ ranging 0.0002 cm/min for raloxifene to 0.004 cm/min for penicillin G and ER_ss_ ranging 0.44 for repaglinide to 4.34 for fluvastatin) were scaled to predict renal clearance (see Methods for details). The predicted renal clearance values spanned 0.006 ml/min/kg for dextromethorphan to 7.4 ml/min/kg for metformin. The MTC predicted renal clearance best for penicillin G, which is predominantly renally cleared. In contrast, the renal clearances of metformin, rosuvastatin, and zidovudine were underpredicted. The IVIVC showed that >30% of the predicted drugs were within a two-fold error.

**Table 2.**
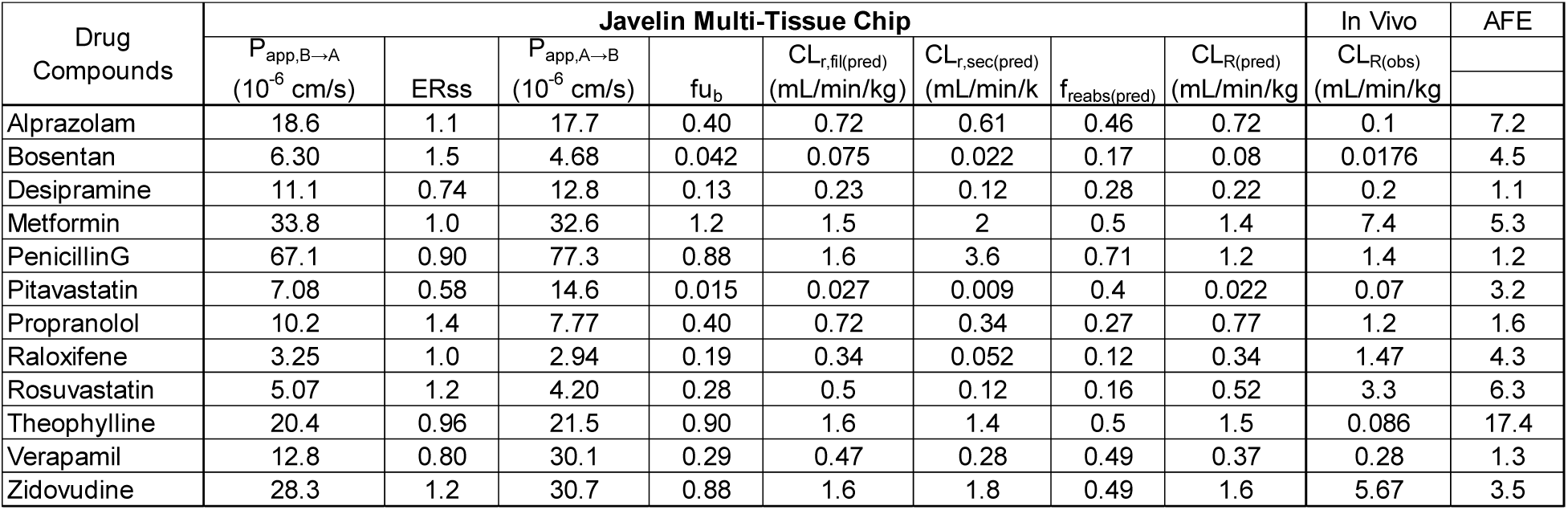
*In vitro in vivo* correlation for renal clearance.

The total clearance (**Fig. 4C**) was estimated by combining the hepatic and renal clearance predictions. The MTC predicted total clearance with >40% of drugs falling within the two-fold error and >60% of drugs falling within the three-fold error.

The volume of distribution (VD_ss_), reflecting the extent of drug distribution in the body, was classified according to acids, bases, or neutrals (**Fig. 4D**). VD_ss_ was predicted using an empirical scaling factor based on the MTC muscle lysate/media ratios spanning 0.014 for rosuvastatin to 2.8 for desipramine (see Methods for details). The corresponding predicted human VD_ss_ value spanned from 0.06 l/kg for rosuvastatin to 12.98 l/kg for desipramine. Comparing MTC predicted VD_ss_ to clinical values showed that >45% of the drugs were predicted within 2-fold and >55% were within three-fold of observed values.

For IVIVE, a PBPK model incorporating both hepatic and renal clearance was adapted to predict clinical concentration profiles from the MTC data (**Fig. 5A**). This method enables prospective prediction of clinical profiles using PBPK models and MTC data without requiring calibration to *in vivo* data. We demonstrate the ability to predict clinical profiles from MTC studies for three drugs: midazolam (hepatically cleared), metformin (renally cleared), and verapamil (mixed hepatic and renal clearance) (**Fig. 5B**). This demonstrates that a single MTC study can be used to predict systemic exposures for drugs with multiple clearance mechanisms.

**Figure 5.**
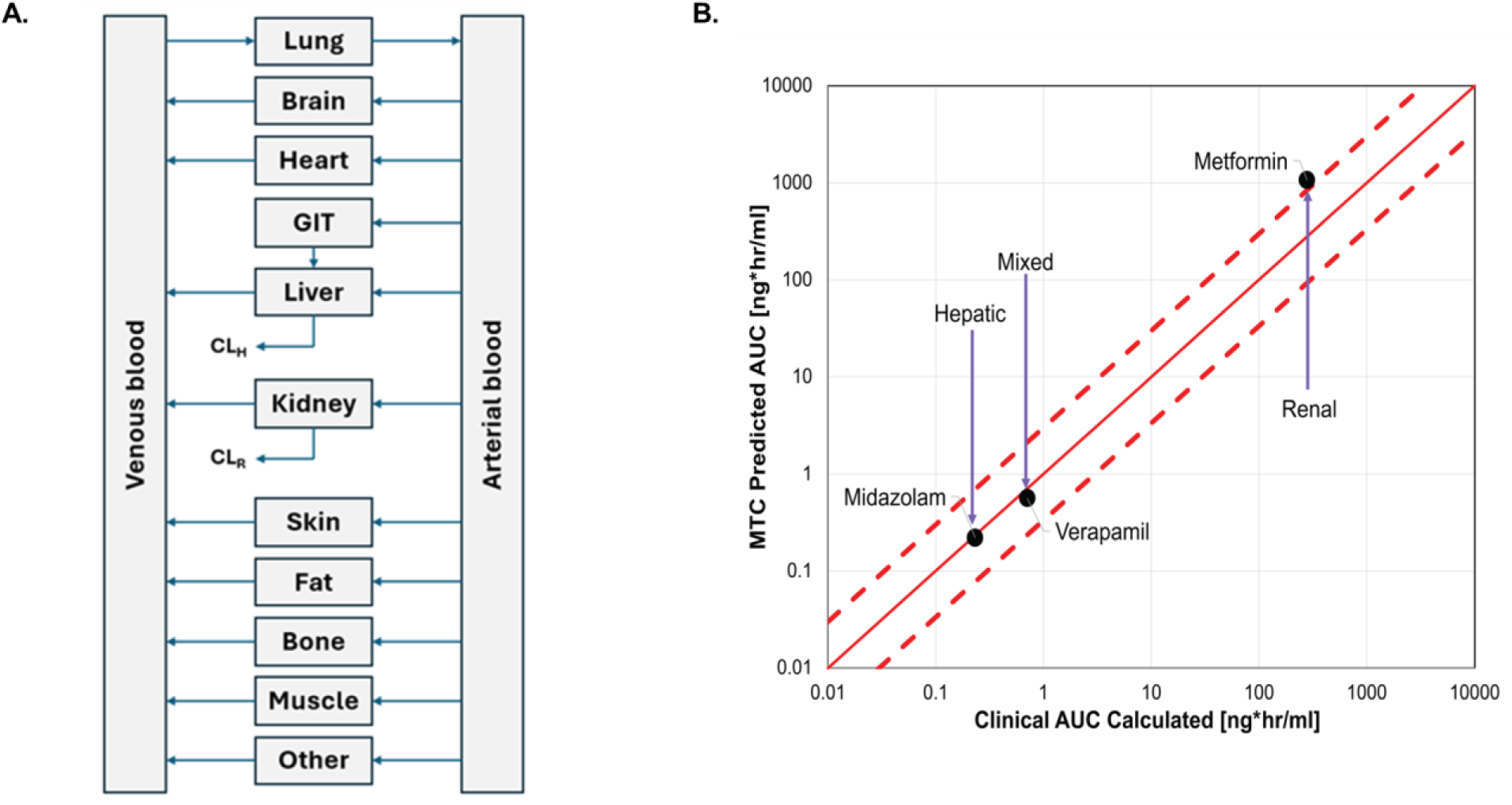
In vitro in vivo extrapolation (IVIVE) and physiologically-based pharmacokinetic (PBPK) modeling for select compounds. (A) Full PBPK model adapted from OSP Model Library by incorporating MTC predicted hepatic clearance (CL_H_) and renal clearance (CL_R_). (B) Drug exposure predicted by MTC-informed PBPK modeling vs. exposures observed in the clinic for three select compounds with different clearance mechanisms. AUC_0-inf_ values were calculated by noncompartmental analysis of the kinetic time-concentration profiles predicted by PBPK modeling or observed in the clinic (see **Supp. Fig. S9** for kinetic profiles). Dashed lines mark three-fold difference between predicted and observed exposures.

## Discussion

The development of the Javelin MTC platform represents a significant step in bridging the gap between *in vitro* and *in vivo* drug testing, particularly in predicting human PK. Traditional animal models and single-tissue chips have limitations in replicating the complex interactions between multiple organs involved in drug ADME (Deng, 2011) (H., 1995) (Martignoni, 2006) (Shanks N, 2009). Our MTC integrates liver, kidney, and skeletal muscle MPSs and offers a more physiologically relevant environment for evaluating drug behavior across multiple organs. This multi-organ system enables the prediction of human PK parameters more accurately than single MPS systems, which typically fail to capture inter-organ interactions that are critical to drug clearance and distribution.

The integration of multiple MPSs within the MTC is crucial for advancing our understanding of ADME processes. By maintaining the long-term functionality and phenotype of each MPS with common medium, the MTC allows for the simultaneous assessment of hepatic, renal, and muscular contributions to drug clearance and disposition. The use of a recirculating media system ensures the viability of the tissues and provides the dynamic conditions necessary for studying complex PK, including both high and low clearance drugs. Importantly, the ability to monitor parent drug depletion through multiple clearance mechanisms and the formation of metabolites in real-time allows for comprehensive PK studies, which is a key advantage over traditional methods.

One of the major advantages of the MTC is its ability to predict a wide range of PK parameters across diverse drugs. As demonstrated in this study, the MTC system accurately predicts hepatic clearance (CL_H_), renal clearance (CL_R_), and volume of distribution (VD_ss_) for multiple drugs, spanning all ECCS classes. This ability to predict drug behavior in both low and high clearance drugs is a major step forward, especially given the challenges associated with predicting the PK of low clearance drugs using current *in vitro* models. The system’s ability to predict these parameters with a three-fold error margin when compared to clinical data confirms the high translational potential of this platform.

Furthermore, the integration of the MTC system with PBPK models delivers accurate clinical predictions. By combining MTC-derived parameters with PBPK modeling, it is possible to extrapolate human clinical profiles from *in vitro* data without relying on *in vivo* animal testing. This process, known as IVIVE, is particularly valuable for the early stages of drug discovery, where preclinical animal studies are costly and time-consuming and often fail to translate to the clinic. This combination of MTC and PBPK modeling has the potential to positively impact drug development by allowing for accurate predictions of clinical PK profiles without using animal experimental data.

The MTC platform’s use of primary human cells is important for maintaining physiological relevance and enabling clinically relevant predictions. The use of primary human hepatocytes, renal cells, and muscle cells ensures that the drug responses observed in the system are reflective of human physiology, reducing the risk of species-specific differences that can arise when using animal-derived cells. Moreover, the use of primary cells supports better prediction of drug metabolism, transport, and potential toxicities. The use of human tissue cells also enhances the system’s ability to study drug-drug interactions (DDIs) in a more realistic context, as shown by the rifampicin- and itraconazole-mediated modulation of CYP3A4 activity in the liver tissue chip (Ohri, Parekh, Nichols, Rajan, & Cirit, 2024).

In the MTC, continuous media recirculation minimizes the issue of nutrient and oxygen depletion in media and helps maintain tissue health over long periods, which is essential for extended drug exposure studies. This setup also enables high-content data generation, which can be used to study drug metabolism, transporter activity, and even biomarkers of disease. The ability to collect samples at multiple time points from both apical and basal compartments provides rich data that can be used to fine-tune understanding of drug disposition in a multi-organ context.

The integration of the MTC system into the drug discovery pipeline has the potential to significantly reduce the need for animal testing, aligning with the FDA Modernization Act 2.0 & 3.0 (S.5002 - 117th Congress (2021-2022): FDA Modernization act 2.0 | congress.gov | library of Congress. (n.d.). https://www.congress.gov/bill/117th-congress/senate-bill/5002; Zushin, 2023; J., 2023) and other regulatory initiatives aimed at reducing animal use in pharmaceutical research. By providing a more accurate and human-specific model for drug testing integrated with PBPK modelling for translation, the MTC platform can help meet the growing demand for alternatives to animal testing while improving the efficiency of drug discovery.

Despite the promising capabilities of the MTC, there are several areas for improvement and expansion. The current version of the MTC platform is optimized for small molecule PK studies but is not yet suited for predicting hepatobiliary clearance, which remains a limitation in the system. The integration of additional systems, such as a gut model for bioavailability studies or lymphatic systems, would further enhance the platform’s versatility and broaden its applicability to other drug modalities, including monoclonal antibodies (mAbs), gene therapies, and cell therapies. Additionally, further developments to the computational workflow could improve MTC performance. For instance, the small molecule PBPK workflow could be expanded to incorporate predicted VDss by adjusted a global Kp scalar to match the MTC-based VDss predictions, and renal clearance estimation could be adjusted to account for the decrease in renal filtrate flow due to water reabsorption (Li, 2020).

Moreover, while the MTC provides valuable insights into drug metabolism and transporter activity, there are areas where more extensive characterization is needed. For instance, the kidney MPS could benefit from additional proteomics and transporter profiling to improve the prediction of renal clearance. Similarly, the introduction of disease models, such as insulin resistance or renal dysfunction, could provide insights into how diseases impact drug metabolism and clearance, which would be invaluable for drug development in therapeutic areas such as diabetes and chronic kidney disease. Additionally, complex immune-competent MPSs are crucial for more accurately modeling disease and assessing ADME properties and immune-mediated toxicities of advanced modalities.

The use of various donor sources for tissue culture could further enhance the robustness and reproducibility of the system, enabling more personalized medicine approaches or capturing population variability. Additionally, incorporating pH gradients in the kidney MPS could improve predictions for acidic drugs, as the current system’s pH levels may not fully replicate the physiological conditions in the renal tubules.

Despite these potential improvements, the MTC platform has already demonstrated substantial advantages over traditional methods in predicting the PK profiles of small molecule drugs. Its ability to model multi-organ interactions, study drug metabolism and transporter activity, and generate high-content data makes it a powerful tool for drug discovery. Further optimization and expansion of the platform will increase its utility in the evaluation of a wider range of drugs, drug interactions, and disease models.

In conclusion, the MTC system represents a promising new approach for drug discovery that can significantly improve the prediction of clinical PK profiles, enhance the study of ADME processes, support the development of new drug modalities and reduce the need for animal experimentation. As the platform continues to evolve, its integration with more complex biological systems and the incorporation of additional technologies such as high-content screening and disease modeling will further cement its role in the future of pharmaceutical research.

