## Supplementary Information for "Predicting Human Pharmacokinetic Parameters of Drugs using a Multi-Tissue Chip Platform Integrating Liver, Kidney, and Skeletal Muscle Microphysiological Systems"

**Supplementary Table 1: Donor Information**

| Age | Race | Gender | BMI |
|---|---|---|---|
| 2 | Hispanic | Male | 14.8 |

**Supplementary Table 2:** Multi-tissue drug cocktails

| Cocktail | Drugs | MW (g/mol) | logP | Ionization |
|---|---|---|---|---|
| Cocktail 1 | Dextromethorphan | 271.40 | 3.5 | Base |
| | Diclofenac | 296.15 | 4.5 | Acid |
| | Midazolam | 325.78 | 3.8 | Neutral |
| | Propranolol | 259.34 | 2.9 | Base |
| Cocktail 2 | Alprazolam | 308.77 | 1.9 | Neutral |
| | Tolbutamide | 270.35 | 2.3 | Acid |
| Cocktail 3 | Bosentan | 551.61 | 4.9 | Acid |
| | S-Warfarin | 308.33 | 3.1 | Acid |
| | Theophylline | 180.16 | -0.3 | Neutral |
| | Zidovudine | 267.24 | -0.3 | Neutral |
| Cocktail 4 | Fluvastatin | 411.46 | 3.8 | Acid |
| | Pitavastatin | 421.47 | 1.9 | Acid |
| | Raloxifene | 473.58 | 4.6 | Base |
| | Verapamil | 454.60 | 4.0 | Base |
| Cocktail 5 | Repaglinide | 452.59 | 4.0 | Acid |
| | Rosuvastatin | 481.54 | 0.9 | Acid |
| Cocktail 6 | Desipramine | 266.38 | 3.9 | Base |
| | Metformin | 129.16 | -2.6 | Base |
| | Penicillin G | 334.40 | 1.5 | Acid |

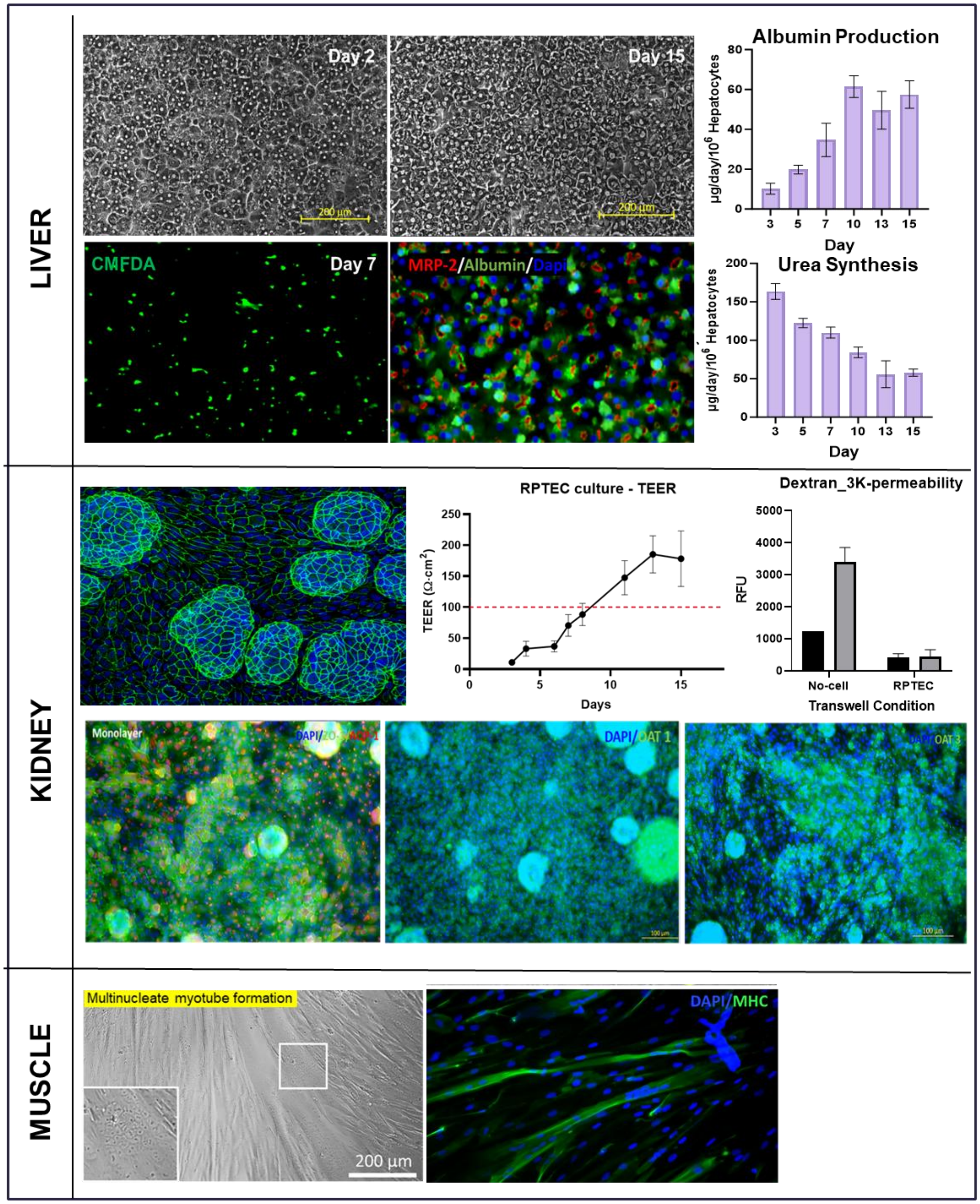


**Supplementary Figure S1. Primary hepatocytes in LTC form physiologically relevant liver tissue** that is evaluated by biochemical assays and Tissue specific protein expression. A) Long-term hepatocyte morphology imaged using bright field microscope on day 2 and 15 showing phenotypic cobblestone morphology. B) Albumin and C) urea levels were assessed in our liver chip system daily on days 4–15 by ELISA and biochemical assay, respectively. B) **Primary human renal proximal tubule epithelial cells (RPTEC) in the kidney tissue chip form an intact barrier that is evaluated by TEER and permeability assays.** A) Brightfield images of RPTECs in Transwell inserts on day 15 with hemicyst formation in between uniform monolayers. B) The barrier integrity of the RPTEC culture was assessed by measuring TEER in inserts and permeability assays using dextran molecule from both apical-basal and basal-apical transport. D) Confocal image of ZO-1-stained tissue. **Primary human skeletal muscle myoblasts form physiologically relevant muscle tissue on Javelin chips.** A) Brightfield image of day 5 myotube culture shows multinucleate myotube formation. C) ICC staining for myosin heavy chain (MHC) expressed in muscle culture at day 5 of maturation.

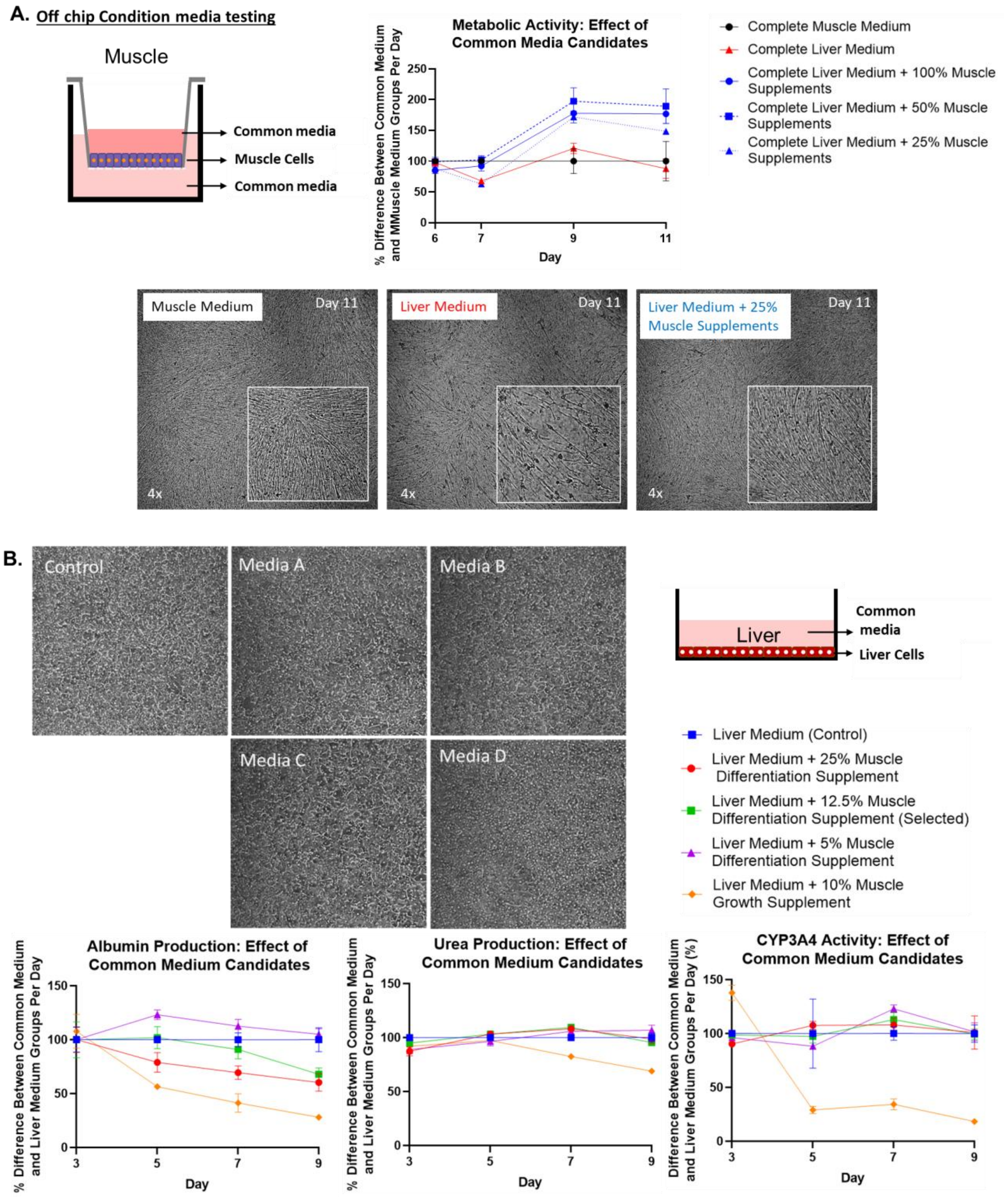

**Supplementary Figure S2. Common media development for multi-MPS integration study.** A) Brightfield image of liver tissue culture after incubating for 3 days under respective media formulation. B) Graph representing normalized effect on different common media formulation on albumin production over 6 days. C) Graph representing the normalized effect of different common media formulations on urea production over 6 days. D) Graph representing normalized effect on different common media formulation on CYP3A4 activity over 6 days.

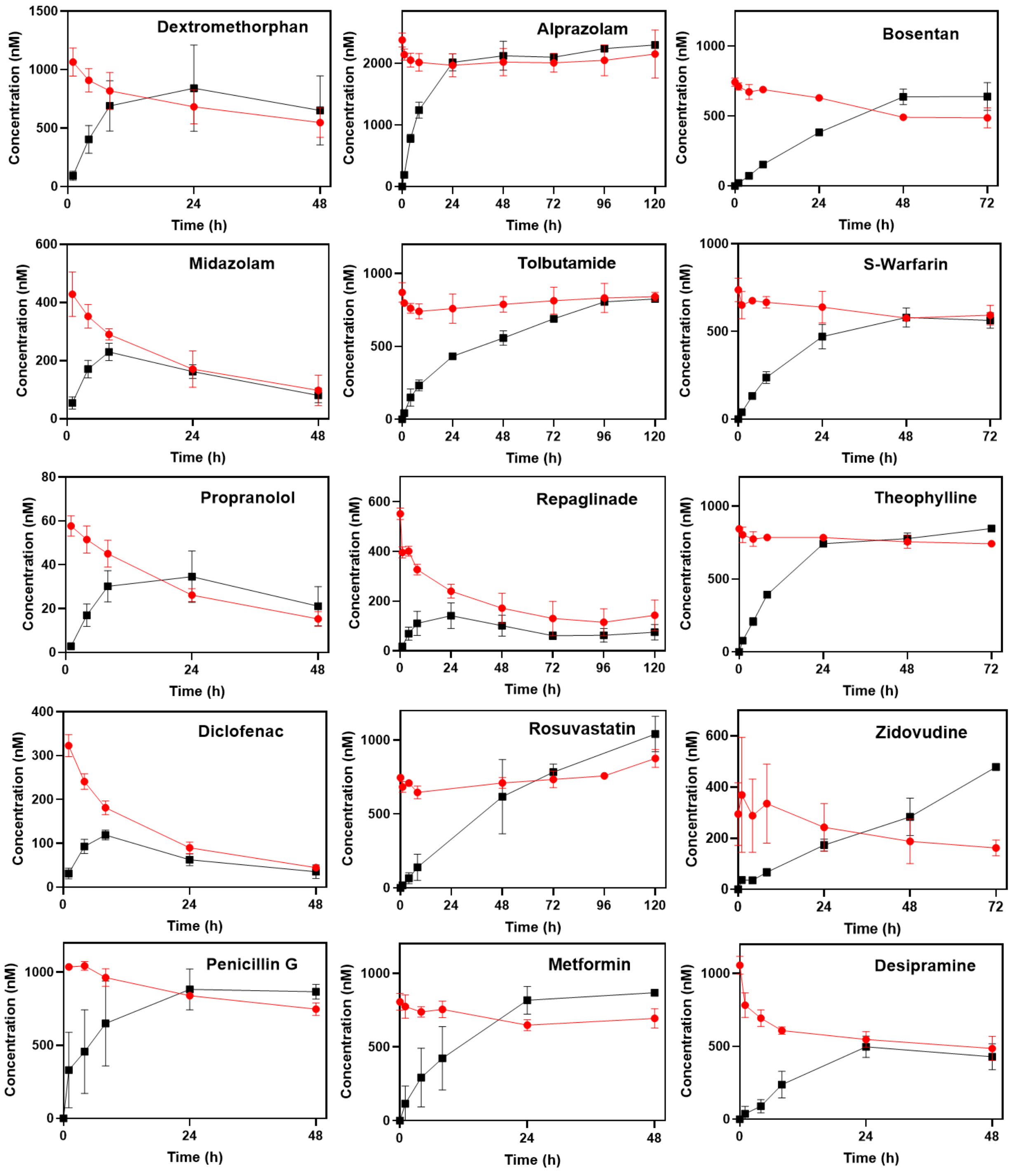

**Supplementary Figure S3**. Time-concentration profiles from ADME studies in the MTC for a diverse set of compounds. Each plot shows the change in drug concentrations observed in the apical kidney compartment (black line) and basal compartment (red lines).

### <u>Supplementary Methods</u>

#### *In vitro data analysis*

This section presents the details of the *in vitro* data analysis to extract on-chip parameters for the liver, kidney, and muscle MPSs.

*Liver MPS analysis*:

Intrinsic clearance was estimated using conservation of mass, noncompartmental analysis, and the inverse well-stirred model.

First, the change in total chip drug concentration over time was calculated for the MTC:

$$C_{MPS}(t) = \frac{C_{basal}(t)V_{basal} + C_{apical}(t)V_{apical}}{V_{basal} + V_{apical}}$$

Where,

$C_{MPS}(t)$: total drug concentration in the MTC at time t.

$C_{basal}(t)$: drug concentration in the basal compartment of the MTC at time t.

$V_{basal}$: volume of the basal compartment in the MTC.

$C_{apical}$: drug concentration in the apical compartment of the MTC at time t.

$V_{apical}$: volume of the apical compartment in the MTC.

Second, a noncompartmental analysis was then applied to $C_{MPS}(t)$ to extract an apparent clearance estimate for the drug in the MTC:

$$CL_{MPS} = \frac{dose}{AUC_{MPS}}$$

Where,

$CL_{MPS}$: Total MPS Clearance

$Dose$: initial amount of drug introduced into the MTC system.

$AUC_{MPS}$: The area under the $C_{MPS}(t)$ curve.

The $AUC_{MPS}$ was computed using the linear up, log down rule on the $n$ observed concentrations and estimated initial concentration, from $C_0$ to $C_n$, and the imputed tail concentrations.

This $CL_{MPS}$ associated with tissue is then applied to an inverse well-stirred model to incorporate recirculation ($Q_{chip}$) and media binding ($fu_{media}$) to derive intrinsic clearance ($CL_{int,u}$) according to:

$$CL_{int,u} = Q_{chip} \cdot \frac{CL_{MPS}}{fu_{media} \cdot (Q_{chip} - CL_{MPS})}$$

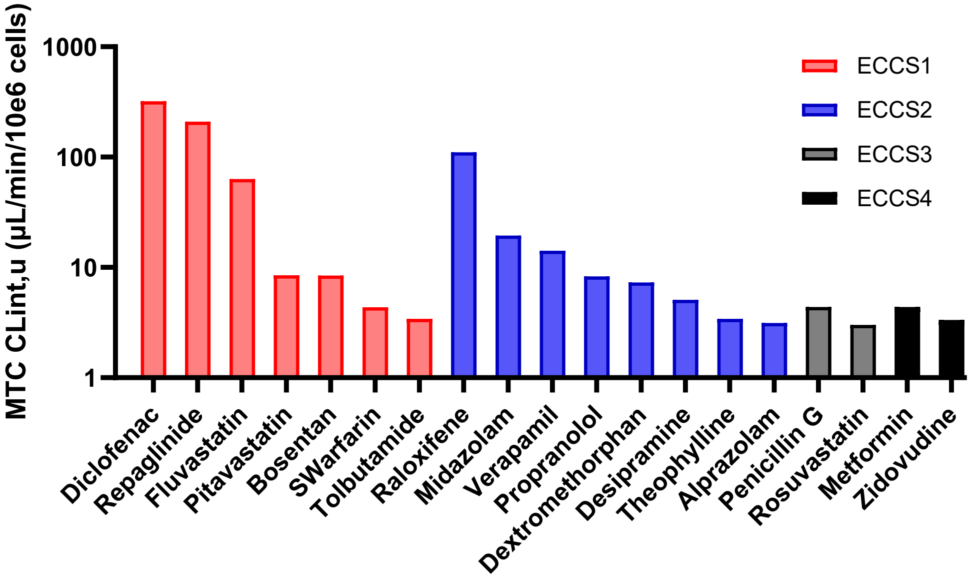


**Supplementary Figure S4**: The calculated $CL_{int,u}$ for the compounds measured in the Multi-Tissue Chip (MTC) system.

*Kidney MPS analysis*:
The kidney analysis involves (1) traditional *in vitro* apparent permeability calculation for $P_{app,\ BA}$ and (2) leveraging the steady-state $ER_{ss}$ observed in the long-term study to estimate $P_{app,\ AB}$ from the same study. $P_{app,\ BàA}$ is estimated from the initial slope of the receiver concentration (i.e., apical kidney) and $ER_{ss}$ from the steady state apical/basal concentration ratio.

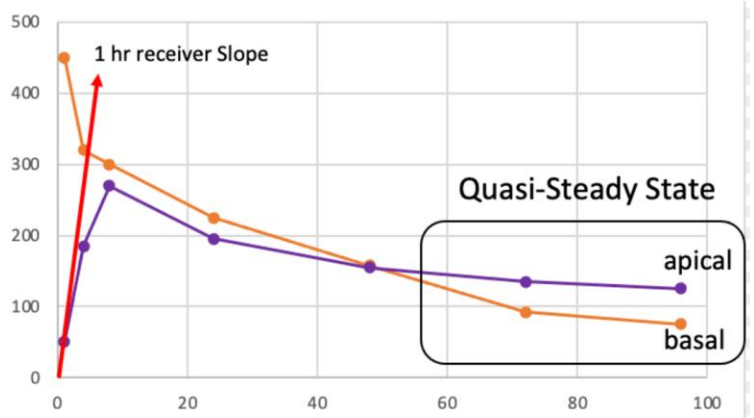


**Supplementary Figure S5:** Time-concentration features used in the kidney MPS analysis: The 1h receiver slope is used in the calculation for $P_{app,\ B\rightarrow A}$, and the quasi-steady-state efflux ratio $(ERss)$ is used in the calculation of $P_{app,\ A\rightarrow B}$. ERss is estimated from the quasi-steady ratio of apical to basal concentrations in the kidney MPS.

1. Given initial basal to apical data, we can calculate $P_{app,B\rightarrow A}$ from slope:

$$P_{app,\,B\to A} = \frac{V_R \cdot C_R}{t \cdot S \cdot C_D} = \frac{V_R}{S \cdot C_D} * (1hr\ slope)$$

where,

$P_{app,\,B\to A}$: Apparent permeability from the basal to apical direction.

$V_R$: Volume of the receiver (apical) compartment.

$C_R$: Concentration of the drug in the receiver compartment.

$t$: Time interval over which permeability is measured.

$S$: Surface area of the membrane separating basal and apical compartments.

$C_D$: Concentration of the drug in the donor (basal) compartment.

2. Given steady state observations (ERss) and $P_{app,B\to A}$, we can estimate $P_{app,A\to B}$:

$$P_{app,A\to B} = \frac{P_{app,B\to A}}{ERss} = \frac{V_R}{S \cdot C_D}\frac{(1hr\ slope)}{ERss}$$

where,

$P_{app,\,A\to B}$: Apparent permeability from the basal to apical direction.

$ER_{ss}$: Steady-state efflux ratio.

*Muscle MPS analysis*:

We have found that the endpoint ratio of lysate to media concentration in the muscle tissue correlates with the steady state volume of distribution. Thus, we quantify for each chip the ratio of lysate/media ratio at the end of each study to predict volume of distribution as described below.

***In vitro* – *in vivo* correlation (IVIVC)**

Predicting clinically-meaningful endpoints from the MTC data involves translating the on-chip parameters calculated above to predicted *in vivo* pharmacokinetic (PK) parameters: hepatic clearance, renal clearance, total clearance, and volume of distribution. The details of each IVIVC calculation will be provided below.

*Hepatic Clearance ($CL_H$):*

The typical value of unbound intrinsic clearance ($\underline{CL}_{int(u)}$) determined for each drug from the pharmacokinetic analysis of the *in vitro* data was scaled up to a human liver equivalent unbound intrinsic clearance ($CL_{int(u),H}$) using

$$CL_{int(u),H} = \underline{CL}_{int(u)} \cdot HC \cdot LW$$

where HC is the human hepatocellularity of 120 million cells / g of liver (Hakooz N 2006) and LW is the average human liver weight of 25.7g / kg of body weight (Brown RP 1997).

The hepatic clearance (referring to whole blood concentrations) was then predicted ($CL_{H(pred)}$) using the Parallel Tube (PT) liver model (Pang KS 1977); for completeness, the hepatic clearance was also calculated using the Well-Stirred (WS) model:

$$CL_{H(pred,PT)} = Q_H \cdot \left(1 - e^{-\cdot fu_b \cdot CL_{int(u),H}/Q_H}\right)$$

$$CL_{H(pred,WS)} = \frac{Q_H \cdot fu_b \cdot CL_{int(u),H}}{Q_H + fu_b \cdot CL_{int(u),H}}$$

where the $Q_H$ is the average hepatic blood flow of 20.7 mL/min/kg of body weight (Davies 1993) and $fu_b$ is the fraction of drug which is unbound in blood.

*Renal Clearance ($CL_R$):*

*In vivo* renal clearance is predicted by

(1) separately estimating glomerular filtration clearance ($CL_{r,fil}$), tubular secretion clearance ($CL_{r,sec}$), and reabsorption fraction ($f_{reab}$), assuming renal metabolic clearance is negligible, and (2) combining the components analytically to estimate total organ clearance.
For the renal clearance analysis, we adapted methods from a kidney transwell system (see Kunze 2014 for details (Kunze 2014)) that are suitable for analysis of data collected from the kidney MPS.

First, intrinsic renal clearance estimates are obtained from apparent permeabilities by scaling with respect to physiology.

$$CL_{r,int,AB/BA} = P_{app,AB/BA} \cdot \frac{\pi \cdot l_{PT} \cdot d_{PT} \cdot n_{neph} \cdot n_{kid}}{BW}$$

where:

$l_{PT}$: length of human proximal tubule (1.5 cm)
$d_{PT}$: diameter of human proximal tubule (7 x $10^{-3}$ cm)
$n_{neph}$: number of nephrons per kidney (1.5 x $10^{6}$)
$n_{kid}$: number of kidneys per human (2)
$BW$: average human body weight (70 kg).

Intrinsic clearances and known physiology are then used to estimate each renal component:

Glomerular filtration clearance:

$$CL_{r,fil} = fu_b \cdot GFR$$

Tubular secretion clearance:

$$CL_{r,sec} = \frac{Q_{r,b} \cdot fu_b \cdot CL_{r,int,\,BA}}{Q_{r,b} + fu_b \cdot CL_{r,int,BA}}$$

Reabsorption fraction:

$$f_{reab} = \frac{CL_{r,int,\,AB}}{GFR + CL_{r,int,AB}}$$

Where:
$fu_b$: Unbound fraction in blood
$GFR$: Glomerular filtration rate (1.79 mL/(min kg))
$Q_{r,b}$: Renal blood flow rate (17.14 mL/(min kg))

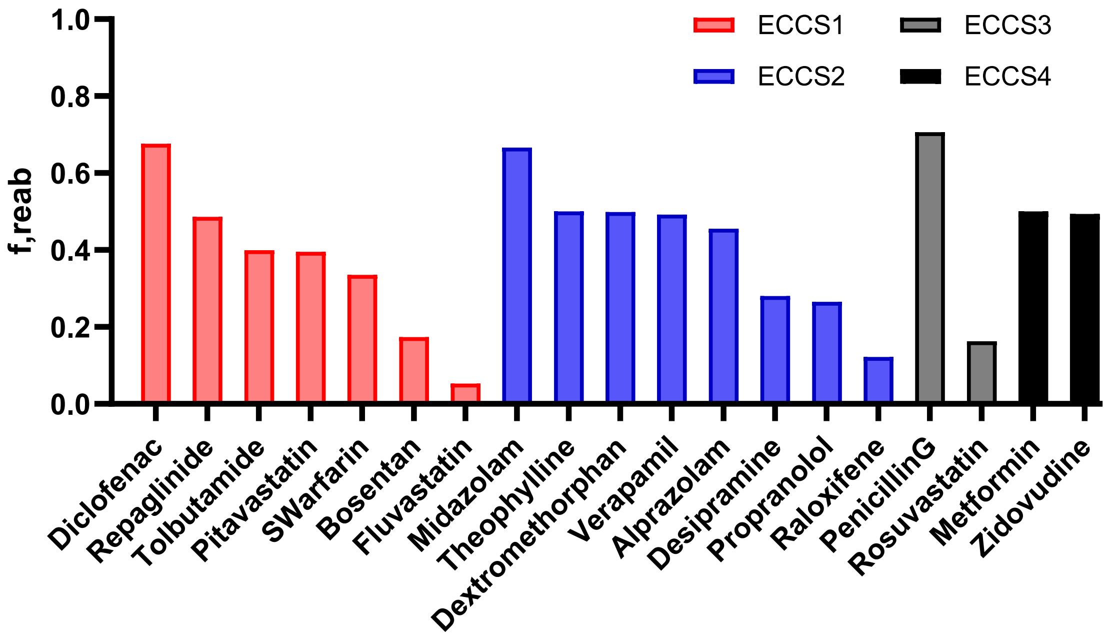


**Supplementary Figure S6:** Reabsorption fraction ($f_{reab}$) for the compounds measured in the Multi-Tissue Chip (MTC) system.

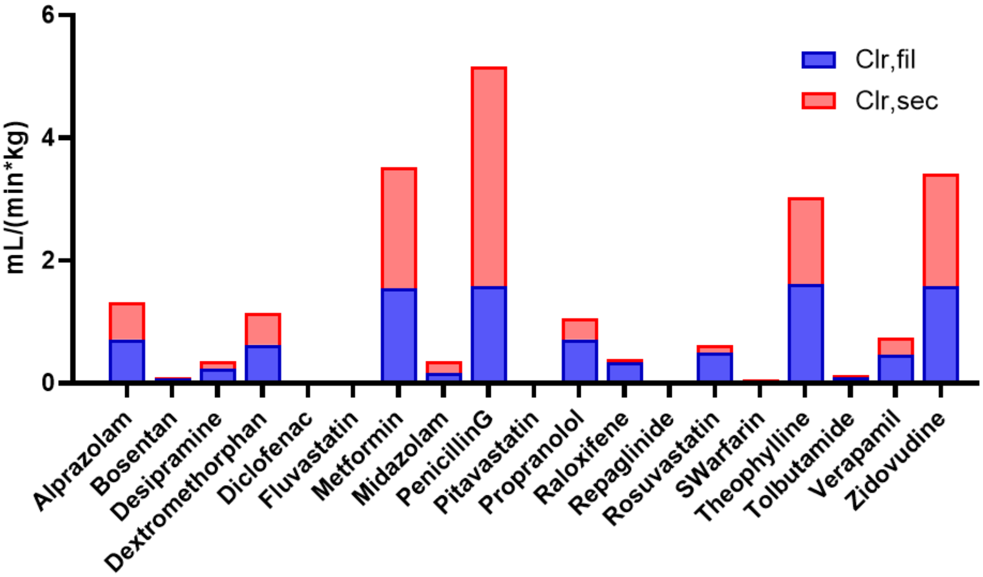


**Supplementary Figure S7:** Tubular secretion clearance ($CL_{r,\mathrm{sec}}$) and Glomerular filtration clearance ($CL_{r,fil}$) for the compounds measured in the Multi-Tissue Chip (MTC) system.

Second, filtration, reabsorption, and secretion are combined to estimate total renal organ clearance:

$$CL_{r,org} = \left(CL_{r,fil} + CL_{r,sec}\right) \cdot (1 - f_{reab})$$

Total Clearance ($CL_{tot}$):
*In vivo* total clearance estimates were then obtained by combining the hepatic and renal clearances:

$$CL_{tot} = CL_{r,org} + CL_H$$

*Steady-state volume of distribution ($VD_{ss}$):*
The steady-state volume of distribution ($VD_{ss}$) was estimated using an empirical scaling factor (ESF) applied to the muscle lysate/media ratio. An ESF = 4.7 was calculated using MTC lysate/media ratios and known $VD_{ss}$ across the full drug set. The same ESF was then used to upscale lysate/media for each drug to predict its clinical $VD_{ss}$.

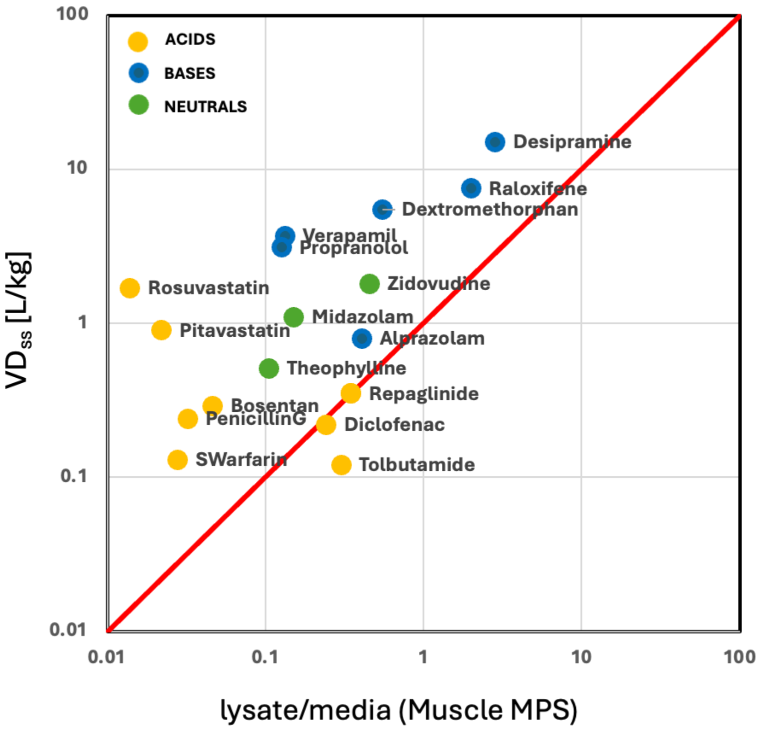


**Supplementary Figure S8:** Estimation of empirical scaling factor with the ratio of lysate-to-media concentration obtained from the muscle MPS to the known clinical volume of distribution ($VD_{ss}$) values.

**<u>In vitro in vivo extrapolation (IVIVE) and PBPK Modeling</u>**
The PBPK model incorporated the standard PK-Sim® model structure comprising 18 compartments, as described by Willmann et al. (Willmann 2003). In this study, built-in compound templates for midazolam (Open-Systems-Pharmacology. (n.d.-b). OSP-PBPK-Model-Library/Midazolam at master · Open-Systems-Pharmacology/OSP-PBPK-Model-Library. GitHub. https://github.com/Open-Systems-Pharmacology/OSP-PBPK-Model-Library/tree/master/Midazolam n.d.), verapamil (Open-Systems-Pharmacology. (n.d.-c). OSP-PBPK-Model-Library/Verapamil at master · Open-Systems-Pharmacology/OSP-PBPK-Model-Library. GitHub. https://github.com/Open-Systems-Pharmacology/OSP-PBPK-Model-Library/tree/master/Verapamil n.d.), and metformin (Open-Systems-Pharmacology. (n.d.). GitHub - Open-Systems-Pharmacology/Metformin-Model: Whole-body PBPK model of metformin

(OCT2/MATE DDI victim drug). GitHub. https://github.com/Open-Systems-Pharmacology/Metformin-Model n.d.) were used as starting points, available in the OSP PBPK model library (Open systems pharmacology. GitHub. (n.d.). https://github.com/Open-Systems-Pharmacology n.d.).

To focus on the experimentally derived PK parameters, we simplified the models by removing default parameters related to specific physiological and enzyme/transporter-mediated mechanisms. Specifically, we deactivated or set to zero the default GFR values and enzyme/transporter activities in PK-Sim® for each compound.

Instead, we incorporated our experimentally predicted plasma clearances directly into the models to capture the net effect of all detailed mechanistic processes.

- Predicted Total Human Hepatic Clearance **(**$CL_{H(pred,PT)}$**)**: The predicted hepatic clearance from the MTC was the input as the plasma clearance in the liver compartment.
- Predicted Total Human Renal Clearance **(**$CL_{r,org}$**)**:The predicted renal clearance from the MTC was the input as the plasma clearance in the kidney compartment.

By integrating our predicted clearance values, we aimed to assess the predictive performance of our Multi-Tissue Chip (MTC)-derived parameters in simulating human pharmacokinetics in PK-Sim®.

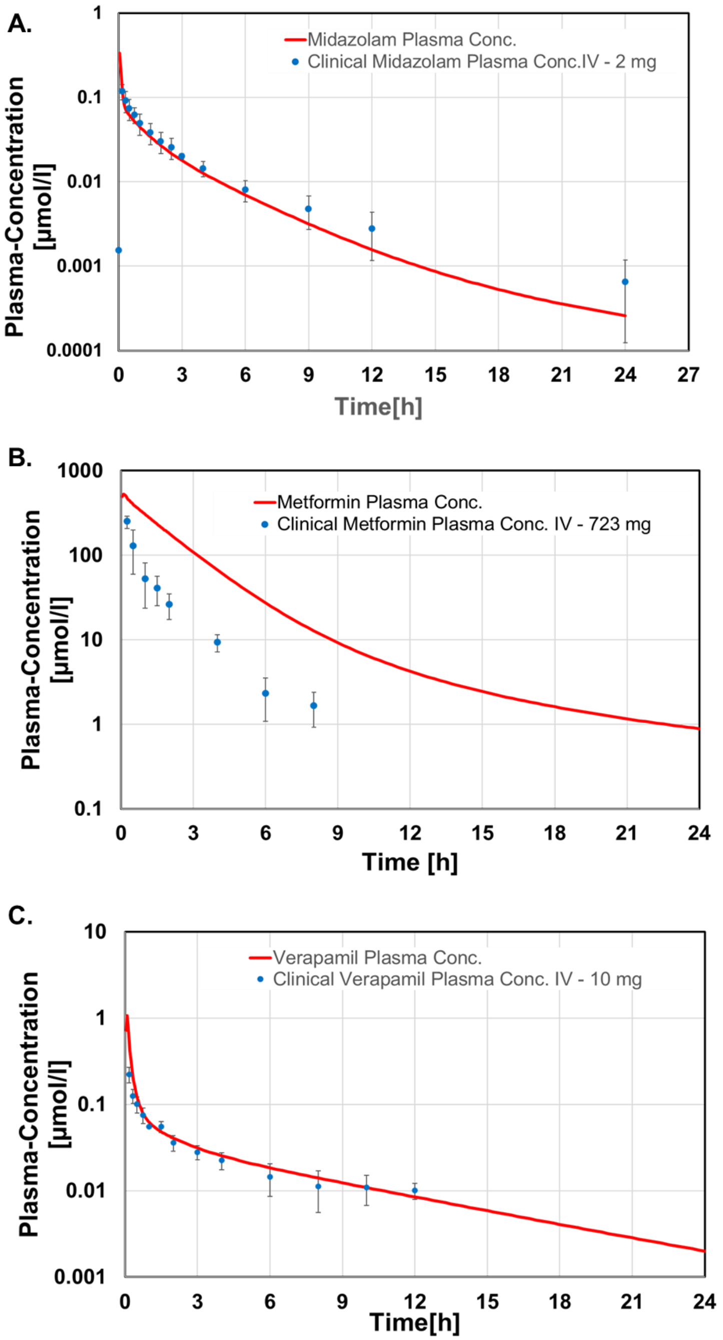


**Supplementary Figure S9**. Kinetic profiles predicted by MTC-informed PBPK modeling (solid lines) and observed in clinic (circles) for three compounds with different clearance mechanisms: (A) midazolam, (B) metformin, (C) verapamil.

**Bioanalysis:** HPLC grade water, HPLC grade water containing 0.1% (v:v) formic acid HPLC grade acetonitrile (ACN), and HPLC grade ACN containing 0.1% (v:v) formic acid were obtained from Fisher Scientific. The following reference material: alprazolam, bosentan, desipramine, dextromethorphan, fluvastatin, midazolam, pitavastatin, propranolol, repaglinide, verapamil, raloxifene, rosuvastatin, S-warfarin, theophylline, zidovudine, diclofenac, metformin, tolbutamide, penicillin G, and furosemide were either provided by Pfizer, Inc. or purchased from Caymen Chemical, Michigan, United States. Stock solutions of each compound were prepared in DMSO or methanol in the range of 1 to 90 mM range. An intermediate solution was prepared in 1:1 acetonitrile: DMSO to contain a concentration of 75,000 nM for each analyte. Standard calibration curves were prepared by serial dilution in HMM – Williams' E medium or HMM + 5% muscle supplement - P/S methanol - cell lysis over the range of 1 to 5000 nM.. Standards were included in the calibration curve if they had a %RE of ≤ 20%. A working internal standard solution was prepared in acetonitrile, containing indomethacin, metoprolol, simvastatin, terfenadine, and zolmitriptan at concentrations of 150, 20.0, 300, 30.0, and 10.0 ng/mL, respectively.

Study samples were prepared by aliquoting 10.0 µL of each sample into a 96-well polypropylene extraction plate. All muscle lysate samples were diluted 10x with HMM + 5% muscle supplement - P/S methanol - cell lysis prior to aliquoting to the 96-well plate extraction plate. Protein precipitation was achieved by the addition of 100 µL to the double blanks and carry over sample and 100 µL of the working internal standard solution to the control blanks and test samples. The 96-well plate polypropylene plate was centrifuged at 1250 g for 5 minutes and 50.0 µL of the supernatant was transferred to a new 96-well polypropylene injection plate. The supernatant was diluted with 200 µL of water containing 0.1% formic acid, with the exception of standards or samples containing metformin, in which 200 µL of 2% heptafluorobutyric (HFBA) acid was added. Metformin is a very polar amine and does not retain on a reverse-phase column. Therefore, 2% HFBA was utilized as a dilution solution to serve as an ion-pairing reagent that creates a neutral ion-pair with metformin, providing retention under reverse-phase chromatography conditions. This ion-pair separates in the mass spectrometer source, allowing the positively charged precursor ion of metformin to enter the mass spectrometer.

The chromatographic system included a Shimadzu Nexera X2 CBM-20A communication bus module, Shimadzu Nexera X2 LC-30AD pumps, Shimadzu Nexera X2 SIL-30ACMP autosampler, and Shimadzu DGU-20A5 degassers. Mobile phase A consisted of water containing 0.1% (v:v) formic acid and mobile phase B consisted of acetonitrile containing 0.1% (v:v) formic acid. The flow rate and column temperatures were 0.5 ml/min and 40°C, respectively. The chromatographic separations were achieved using a BEH C18 column (2.1 mm x 50 mm, 1.7 μm, P/N 186002350). 10 . The chromatographic gradient is described in Supplementary Table 3.

**Supplementary Table 3:** Chromatographic Gradient

| Time | Module | Evemt | Parameter |
|---|---|---|---|
| 0.01 | Pumps | Pump B Conc. | 2 |
| 0.70 | Pumps | Pump B Conc. | 2 |
| 2.10 | Pumps | Pump B Conc. | 95 |
| 2.60 | Pumps | Pump B Conc. | 95 |
| 2.70 | Pumps | Pump B Conc. | 2 |
| 4.00 | Controller | Stop | |

The mass spectrometric detector utilized to conduct sample analysis was a Sciex Triple Quad™ 6500(+) equipped with a IonDrive™ Turbo V source. The source conditions were as follows: positive ion electrospray, CUR 20, CAD 8, IS 4500, TEM 600, GS1 50, GS2 60, EP 10, and CXP The data acquisition was performed using multiple reaction monitoring (MRM). MRM transitions and mass spectrometric parameters can be found in Supplementary Table 4**.** Data acquisition and chromatographic review were performed using Applied Biosystems/MDS Sciex Analyst, version 1.7.2. The calibration curves utilized $1/x^2$ weighting.

**Supplementary Table 4:** Mass Spectrometric Parameters

| Analytes | | | | | | | |
|---|---|---|---|---|---|---|---|
| **Precursor Ion** | **Product Ion** | **Dwell time** | **Compound** | **DP** | **CE** | **RT** | **IS** |
| 309.2 | 205.2 | 15.0 | Alprazolam | 70.0 | 60.0 | 2.22 | Terfenadine |
| 553.0 | 202.0 | 15.0 | Bosentan | 58.0 | 42.0 | 2.37 | Motoprolol |
| 267.3 | 193.2 | 15.0 | Desipramine | 30.0 | 55.0 | 2.04 | Terfenadine |
| 272.3 | 215.2 | 15.0 | Dextramethorphan | 80.0 | 35.0 | 1.98 | Metoprolol |
| 411.9 | 224.0 | 15.0 | Fluvastatin | 15.0 | 30.0 | 2.40 | Indomethacin |
| 326.1 | 222.2 | 15.0 | Midazolam | 60.0 | 60.0 | 1.99 | Metoprolol |
| 421.9 | 290.1 | 15.0 | Pitavastatin | 60.0 | 42.0 | 2.11 | Terfenadine |
| 260.2 | 116.2 | 15.0 | Propranolol | 80.0 | 30.0 | 1.96 | Metoprolol |
| 453.2 | 162.4 | 15.0 | Repaglinide | 55.0 | 30.0 | 2.22 | Terfenadine |
| 455.2 | 165.4 | 15.0 | Verapamil | 60.0 | 45.0 | 2.06 | Terfenadine |
| 474.3 | 112.1 | 15.0 | Raloxifene | 122.0 | 44.0 | 1.97 | Metoprolol |
| 482.1 | 258.1 | 15.0 | Rosuvastatin | 140.0 | 40.0 | 2.23 | Terfenadine |
| 309.1 | 251.1 | 15.0 | S-Warfarin | 20.0 | 25.0 | 2.34 | Metoprolol |
| 181.1 | 124.2 | 15.0 | Theophylline | 20.0 | 15.0 | 1.82 | Zolmitriptan |
| 268.1 | 127.0 | 15.0 | Zidovudine | 20.0 | 15.0 | 1.82 | Zolmitriptan |
| 296.1 | 214.0 | 15.0 | Diclofenac | 70.0 | 40.0 | 2.41 | Metoprolol |
| 130.1 | 60.1 | 15.0 | Metformin | 45.0 | 25.0 | 0.99 | Zolmitriptan |
| 271.0 | 172.0 | 15.0 | Tolbutamide | 66.0 | 18.0 | 2.26 | Terfenadine |
| 335.14 | 160.11 | 15.0 | Penicillin G | 40.0 | 25.0 | 1.89 | Terfenadine |
| 329.19 | 284.94 | 15.0 | Furosamide | -55.0 | -25 | 2.15 | No internal Standard used. |

| Internal Standards | | | | | | |
|---|---|---|---|---|---|---|
| **Precursor Ion** | **Product Ion** | **Dwell time** | **Compound** | **DP** | **CE** | **RT** |
| 472.1 | 436.0 | 15.0 | Terfenadine | 80.0 | 30.0 | 2.16 |
| 358.3 | 139.2 | 15.0 | Indomethacin | 46.0 | 26.0 | 2.41 |
| 268.1 | 116.1 | 15.0 | Metoprolol | 60.0 | 25.0 | 1.87 |
| 288.2 | 243.2 | 15.0 | Zolmitriptan | 65.0 | 25.0 | 1.71 |

n.d. "Open systems pharmacology. GitHub. (n.d.). https://github.com/Open-Systems-Pharmacology ."

n.d. "Open-Systems-Pharmacology. (n.d.). GitHub - Open-Systems-Pharmacology/Metformin-Model: Whole-body PBPK model of metformin (OCT2/MATE DDI victim drug). GitHub. https://github.com/Open-Systems-Pharmacology/Metformin-Model."

n.d. "Open-Systems-Pharmacology. (n.d.-b). OSP-PBPK-Model-Library/Midazolam at master · Open-Systems-Pharmacology/OSP-PBPK-Model-Library. GitHub. https://github.com/Open-Systems-Pharmacology/OSP-PBPK-Model-Library/tree/master/Midazolam."

n.d. "Open-Systems-Pharmacology. (n.d.-c). OSP-PBPK-Model-Library/Verapamil at master · Open-Systems-Pharmacology/OSP-PBPK-Model-Library. GitHub. https://github.com/Open-Systems-Pharmacology/OSP-PBPK-Model-Library/tree/master/Verapamil."

Pang KS, Rowland M. 1977. "Hepatic clearance of drugs. I. Theoretical considerations of a "well-stirred" model and a "parallel tube" model. Influence of hepatic blood flow, plasma and blood cell binding, and the hepatocellular enzymatic activity on hepatic drug clearance." *Journal of pharmacokinetics and biopharmaceutics* 5 (6): 625–653.

Willmann, S., Lippert, J., Sevestre, M., Solodenko, J., Fois, F., & Schmitt, W. 2003. "PK-Sim®: a physiologically based pharmacokinetic 'whole-body'model." *Biosilico* 1 (4): 121-124.
